# Wnt/β-catenin signalling underpins juvenile *Fasciola hepatica* growth and development

**DOI:** 10.1101/2024.09.04.611166

**Authors:** Rebecca Armstrong, Nikki J Marks, Timothy G Geary, John Harrington, Paul M Selzer, Aaron G Maule

**Affiliations:** Understanding Health & Disease, School of Biological Sciences, Queen’s University Belfast, Belfast BT9 5DL, UK; Institute of Parasitology, McGill University, Ste-Anne-de-Bellevue, QC H9X 3V9, Canada; Boehringer Ingelheim Animal Health, Duluth, GA 30096-4640, USA; Boehringer Ingelheim Vetmedica GmbH, Binger Str. 173 | 55216 Ingelheim am Rhein, Germany

**Keywords:** Fasciola, development, Wnt, neoblast, RNAi, FISH

## Abstract

Infection by the liver fluke, *Fasciola hepatica*, places a substantial burden on the global agri-food industry and poses a significant threat to human health in endemic regions. Widespread resistance to a limited arsenal of chemotherapeutics, including the frontline flukicide triclabendazole (TCBZ), renders *F. hepatica* control unsustainable and accentuates the need for novel therapeutic target discovery. A key facet of *F. hepatica* biology is a population of specialised stem cells which drive growth and development - their dysregulation is hypothesised to represent an appealing avenue for control. The exploitation of this system as a therapeutic target is impeded by a lack of understanding of the molecular mechanisms underpinning *F. hepatica* growth and development. Wnt signalling pathways govern a myriad of stem cell processes during embryogenesis and drive tumorigenesis in adult tissues. Here, we identify five putative Wnt ligands and five Frizzled receptors in liver fluke transcriptomic datasets and find that Wnt/β-catenin signalling is most active in juveniles, the most pathogenic life stage. FISH-mediated transcript localisation revealed partitioning of the five Wnt ligands, with each displaying a distinct expression pattern, consistent with each Wnt regulating the development of different cell/tissue types. The silencing of each individual Wnt or Frizzled gene yielded significant reductions in juvenile worm growth and, in select cases, blunted the proliferation of neoblast-like cells. Notably, silencing *Fh*CTNNB1, the key effector of the Wnt/β-catenin signal cascade led to aberrant development of the neuromuscular system which ultimately proved fatal - the first report of a lethal RNAi-induced phenotype in *F. hepatica*. The absence of any discernible phenotypes following the silencing of the inhibitory Wnt/β-catenin destruction complex components is consistent with low destruction complex activity in rapidly developing juvenile worms, corroborates transcriptomic expression profiles and underscores the importance of Wnt signalling as a key molecular driver of growth and development in early stage juvenile fluke. The pharmacological inhibition of Wnt/β-catenin signalling using commercially available inhibitors phenocopied RNAi results and provides impetus for drug repurposing. Taken together, these data functionally and chemically validate the targeting of Wnt signalling as a novel strategy to undermine the pathogenicity of juvenile *F. hepatica*.

**AUTHOR SUMMARY:** The liver fluke, *Fasciola hepatica* significantly undermines the health and welfare of livestock worldwide and causes fascioliasis, a neglected tropical disease of humans. The most damaging stage of liver fluke infection is caused by the migration of juvenile worms within the liver tissue. Of all drugs approved for liver fluke treatment, just one, triclabendazole (TCBZ), is active on this pathogenic juvenile stage. TCBZ resistance is now widespread rendering liver fluke control unsustainable. This highlights the need for novel drug target identification and validation. A key aspect of juvenile worm biology is their ability to rapidly grow and develop, processes driven by a population of specialised stem cells. As such, the dysregulation of stem cells represents an attractive avenue for liver fluke control. One molecular pathway known to regulate stem cell dynamics in higher organisms is the Wnt signalling pathway. Bioinformatic searches of gene sequence datasets identified all major signalling components of both canonical and non-canonical Wnt pathways in *F. hepatica*. The localisation of *Fh*Wnt pathway components revealed remarkably distinct and widespread expression patterns throughout the *F. hepatica* body. Gene silencing of putative *Fh*Wnt pathway components revealed that those involved in the Wnt/β-catenin signal cascade are fundamental to juvenile growth and, in some cases, stem-like cell proliferation. The silencing of liver fluke β-catenin led to aberrant neuromuscular development and proved lethal to juvenile fluke. Biweekly exposures to commercially available Wnt pathway inhibitory compounds phenocopied the delayed development observed in the gene silencing experiments. These data suggest that *Fh*Wnt pathway components represent attractive targets for the development of novel flukicides or indeed, the repurposing of existing Wnt antagonists for parasite control.

## INTRODUCTION

The liver fluke, *Fasciola hepatica* is an aetiological agent of fasciolosis, a disease that significantly undermines the health and productivity of ruminant livestock worldwide and costs the global agri-food industry in excess of US$3.2 billion annually [1]. The ability of this pervasive organism to transcend host species barriers has also rendered *F. hepatica* a significant threat to human health, such that fascioliasis is recognised by the World Health Organisation as both a Neglected Tropical Disease and an emerging zoonosis [2]. Control is precarious, depending heavily upon a limited arsenal of drugs, notably triclabendazole (TCBZ), unique amongst the flukicides in its ability to treat both acute and chronic forms of fasciolosis [3]. Heavy overreliance on TCBZ has inevitably selected for resistance, cases of which have been reported in both animal and human hosts [4, 5, 6], accentuating the pressing need for novel therapeutic target discovery.

Wnt signal transduction pathways comprise tightly concerted chains of molecular events that execute pivotal roles during embryogenesis and adult tissue homeostasis by orchestrating a panoply of cellular processes, including proliferation, differentiation, migration and self-renewal [7, 8, 9]. Accordingly, Wnt proteins serve as directional growth factors, facilitating precision morphogenesis and axial patterning during development. The initiation of the Wnt signal cascade is contingent upon the secretion of Wnt proteins into the extracellular matrix and their subsequent interaction with frizzled (FZD) G-protein coupled receptors (GPCRs) on the cell surface. In mammals, the Wnt family of ligands is represented by 19 40kDa glycoproteins, each with 23-24 conserved, invariantly spaced cysteine residues. Traditionally, Wnt signalling is segregated into canonical and non-canonical branches, such that dependent upon which they initiate, Wnts too are designated either canonical or non-canonical. As may be inferred from its fundamental role in such a myriad of biological processes, Wnt signal transduction is inherently complex and more integrated than originally believed, with Wnt-FZD relationships demonstrating a high degree of promiscuity. Nevertheless, the classical canonical and non-canonical nomenclature remains convenient with regards to discussion.

The prototype Wnt pathway, presently designated the canonical or Wnt/β-catenin pathway, was first described in the early 1980s in *Drosophila melanogaster*, with the discovery that mutations in the wingless (Wg) gene produced flies devoid of halteres. Parallel research identified the murine proto-oncogene Int-1 as a driver of mammary tumorigenesis in mice, however, it was not until 1987 that Int-1 and Wg were deemed orthologous, giving rise to the portmanteau, ‘Wnt’ [10].

Wnt pathway orthologues have been identified in a range of organisms [11], including those belonging to the most basal animal phylum, the Porifera [12], highlighting Wnt signalling as a true innovation of metazoan evolution. Indeed, the significance of this striking conservation in platyhelminth development was recently exposed by the discovery that Wnt/β-catenin signalling governs head/tail specification during planarian regeneration [13, 14, 15]. Further, reports of diverse, aberrant phenotypes arising from RNAi-mediated silencing of various Wnt pathway orthologues in planaria is indicative of an exceptionally intricate regulatory network fundamental to platyhelminth biology [16, 17, 18].

Unsurprisingly, homologues of Wnt pathway components have also been identified in the genomes of parasitic species including *Hymenolepis microstoma, Echinococcus spp, Schistosoma spp,* and *F. hepatica* [19, 20, 21]. Comparative expression analyses of Wnt proteins and antagonists in *E. multilocularis* and *H*. *microstoma*, revealed that just as in planaria, Wnt signalling is a key determinant of anterior/posterior (A/P) specification in cestodes [22]. The isolation and immunolocalisation of *S. japonicum Sj*WNT5 and *Sj*FZD7 demonstrated high levels of expression in the schistosomula stages and immunoreactivity within the musculature and reproductive apparatus [23, 24], suggestive of roles in muscle development. Additionally, recent transcriptomic analyses in *F. hepatica* have alluded to the significance of Wnt signalling in the growth and development of juvenile stage fluke [25, 26]. Although such studies support the development of hypotheses on the possible roles and relative importance of selected Wnt pathway components in the biology of these species, data pertaining to the functional interrogation of Wnt signalling and indeed, developmental pathways in parasitic groups, remain scant.

Aberrant Wnt signalling has long been pathophysiologically linked to a plethora of pathologies, including neurodegeneration, birth defects and carcinogenesis. The role of Wnt in carcinogenesis is most comprehensively described in relation to the maintenance of cancer stem cell (CSC) ‘stemness’ in colorectal cancer, however, mutations in the expression of various Wnt/β-catenin pathway components also drive the development and metastasis of many other cancers [27, 28, 29]. As such, the pharmacological targeting of Wnt pathways is the subject of intensive research in the pursuit of novel cancer therapeutics.

Parasite survival, propagation and virulence is contingent upon their growth and development *in vivo*. In parasitic platyhelminths, these are processes driven by the proliferation of specialised stem cells. For example, the germinative cells of *Echinococcus spp* drive metacestode growth and persist following benzimidazole treatment, facilitating the recurrence of disease [30, 31]. Additionally, the neoblast-like cells of *Schistosoma spp* drive tegumental renewal at the host-parasite interface, thereby contributing to longevity [32]. While the significance of stem cell-driven growth and development in parasites is evident, the molecular mechanisms underpinning their dynamics have yet to be elucidated. Given the vital role of Wnt signalling in governing the growth, development and stem cell dynamics of higher organisms, it seems reasonable to hypothesise that interfering with Wnt signalling in juvenile *F. hepatica* would undermine growth and development, such that it may constitute an attractive source of novel flukicide targets. Moreover, the availability of a large pool of Wnt pathway inhibitors and agonists provides impetus for compound screens in helminth parasites, with the aim of identifying those suitable for ‘repurposing’ as novel anthelmintics.

Motivated by the availability of *F. hepatica* ‘omics’ datasets and the amenability of *in vitro* maintained juvenile fluke to RNAi over extended time periods, this study aimed to characterise and functionally interrogate *Fh*Wnt pathway components to uncover potentially druggable targets integral to the growth and development of juvenile *F. hepatica*.

### RESULTS & DISCUSSION

Increasing reports of resistance to the frontline flukicide, TCBZ accentuates the urgent need for novel therapeutic target discovery and validation in the liver fluke. The recent introduction of *F. hepatica* into the ‘omics’ era, coupled with a robust RNAi methodology provides impetus for functional genomic studies to elucidate gene function. Moreover, the development of an *in vitro* culture platform, permitting long-term survival *in vitro* and supporting the proliferation of neoblast-like cells and growth allows, for the first time, the interrogation of *F. hepatica* growth and development on a molecular level. This study exploited the *F. hepatica* molecular toolbox to annotate and functionally probe evolutionarily conserved Wnt signal transduction pathways in juvenile *F. hepatica,* with the aim of uncovering genes integral to growth, development and neoblast-like cell dynamics.

#### *Fasciola hepatica* possess the core components of functional Wnt signalling pathways

Mining genomic datasets facilitated the identification of putative orthologues of all major canonical and non-canonical Wnt pathway components in *F. hepatica*, including receptors and antagonists. Previously, McVeigh *et al.* [21] reported a greatly reduced and dispersed complement of Wnt and Frizzled orthologues in *F. hepatica*, comprising three putative Wnt protein ligands and five Frizzled class GPCRs. Here, this was corroborated by leveraging the most recent *F. hepatica* genome assemblies (PRJEB25283 and PRJNA179522), with sequence annotations revealing *Fh*Wnts to be representative of the WNT1, WNT4 and WNT9 subfamilies. Two additional Wnt ligands belonging to the WNT2 and WNT5 subfamilies were also identified. All gene annotations were assigned based upon the top hit generated from reciprocal BLASTs against *H. sapiens* using the NCBI non redundant protein (nr) database. While this represents a greatly reduced complement of both protein families when compared to mammalian genomes which typically harbour 19 Wnts and 10 FZDs, similar numbers have been reported in other members of the phylum. For example, the genome of *S. mediterranea* encodes 9 Wnts and 13 FZDs, *S. mansoni* 5 Wnts and 9 FZDs, while *Echinococcus spp* have 6 Wnts and 8 FZDs [19]. Both *F. hepatica* and *S. mansoni* were found to lack orthologues of low-density lipoprotein receptor-related proteins 5 and 6. This constitutes a noteworthy absence given the key roles of LRP5/6 as co-receptors in the Wnt/β-catenin pathway. Further, while the related *S. mediterranea* and *E. multilocularis* lack LRP5, they do possess LRP6 orthologues. In the absence of LRP5/6, it may be that *F. hepatica* and *S. mansoni* instead opt to employ receptor tyrosine kinase like orphan receptors 1 and 2 (ROR1/ROR2) as co-receptors in their respective Wnt/β-catenin signal cascades and as such, these proteins warrant future functional interrogation. Additionally, *F. hepatica* possess a LRP1B orthologue (FhHiC23_g15024). LRP1B is not recognised as a Wnt/β-catenin pathway component in higher organisms, however, there exist reports of interactions with Wnt and FZDs [44, 45]. While the results of such interactions repress Wnt signalling, it suggests that the docking site of LRP1B is structurally and functionally compatible with Wnt protein ligands and as such, *Fh*LRP1B may have displaced *Fh*LRP5/6 over the course of evolution. While the core molecular machinery required for functional Wnt pathways is clearly evident in *F. hepatica*, it is noteworthy that much like other parasitic flatworm species, they lack homologues of the O-acyltransferase porcupine (PORCN) and the antagonists Dikkopf and Cerberus. A third antagonist, Wnt inhibitory factor (WIF), was not identified in liver fluke datasets, despite its presence in the related *Hymenolepis microstoma, Echinococcus spp, Schistosoma spp* and *S. mediterranea* [19]. While two *Fh*SFRPs were identified, there is a general paucity of pathway antagonists within the *F. hepatica* genome, a pattern which appears to be unanimous across several basal metazoan lineages, including *Drosophila* and *C. elegans* [46, 47].

With few exceptions, a full complement of orthologues unique to both non-canonical pathways were also identified (S2 Table), with the conserved domains of each gene family intact, indicating that both canonical and non-canonical Wnt signalling is functional in *F. hepatica*. The sole planar cell polarity (PCP) pathway component lacking any discernible orthologue in *F. hepatica* was dishevelled-associated activator of morphogenesis 1 (DAAM1), while nuclear factor of activated T-cells (NFAT) and cell division control protein 42 homolog (CDC42) were absent from the *Fh*Wnt/Ca^2+^ pathway.

#### *Fh*Wnt pathway orthologues possess key domains associated with gene function in higher organisms

Of the three characterised Wnt pathways, the Wnt/β-catenin signal cascade is the most widely studied and its significance in platyhelminth development was recently underscored in regenerating *Planaria*. As such, putative *Fh*Wnt/β-catenin pathway orthologues were subjected to further *in silico* analyses to validate their integrity as true orthologues of those annotated in both higher and related organisms. The predicted protein sequences of all genes tentatively selected for RNAi experiments were aligned with corresponding sequences from *H. sapiens*, *M. musculus, D. melanogaster, S. mediterranea, E. multilocularis* and *S. mansoni*.

Alignments of *Fh*Wnts revealed full conservation of the Wnt family signature motif (CKCHGVSGSC) within the frizzled receptor binding site [33], while the conserved C-terminal cytoplasmic motif of frizzled receptors, (KTXXXW, where X denotes any amino acid), required for both the activation of the Wnt/β-catenin pathway, and for membrane localisation and phosphorylation of dishevelled [34], is present in *Fh*FZDs (Fig. 1A & 1B). *Fh*CTNNB1 possesses N-terminal residues essential for protein-level regulation including Ser33/Ser37/Thr41 and Ser45 which are phosphorylated by GSK3β and CK1 of the destruction complex, respectively. Also in consensus with higher and related organisms, was a conserved lysine residue at position 49 and the DSG motif (DSGοXS where ο denotes a hydrophobic amino acid, and X any amino acid), in which both serines are phosphorylated and recognised by the E3 ubiquitin ligase complex, resulting in the ubiquitination and degradation of β-catenin [35] (Fig. 1D). *Fh*DSHs were found to possess the characteristic dishevelled PDZ binding domain [36, 37] in addition to the class III PDZ-binding motif [38] (Fig. 1C). The highly conserved motifs CDFGSAK and SYICSR [39], present only within members of the GSK-3 subfamily of serine/threonine protein kinases were identified in both *Fh*GSK3β isoforms (Fig. 1F). *Fh*APC possesses two SAMP repeats, believed to mediate interactions with axin of the destruction complex [40] (Fig. 1E). Finally, alignments of the antagonists, *Fh*SFRPs revealed the presence of a fully conserved cysteine rich domain homologous to that of FZDs, containing 10 invariantly spaced cysteine residues [41] (Fig. 1G).

**Figure 1.**
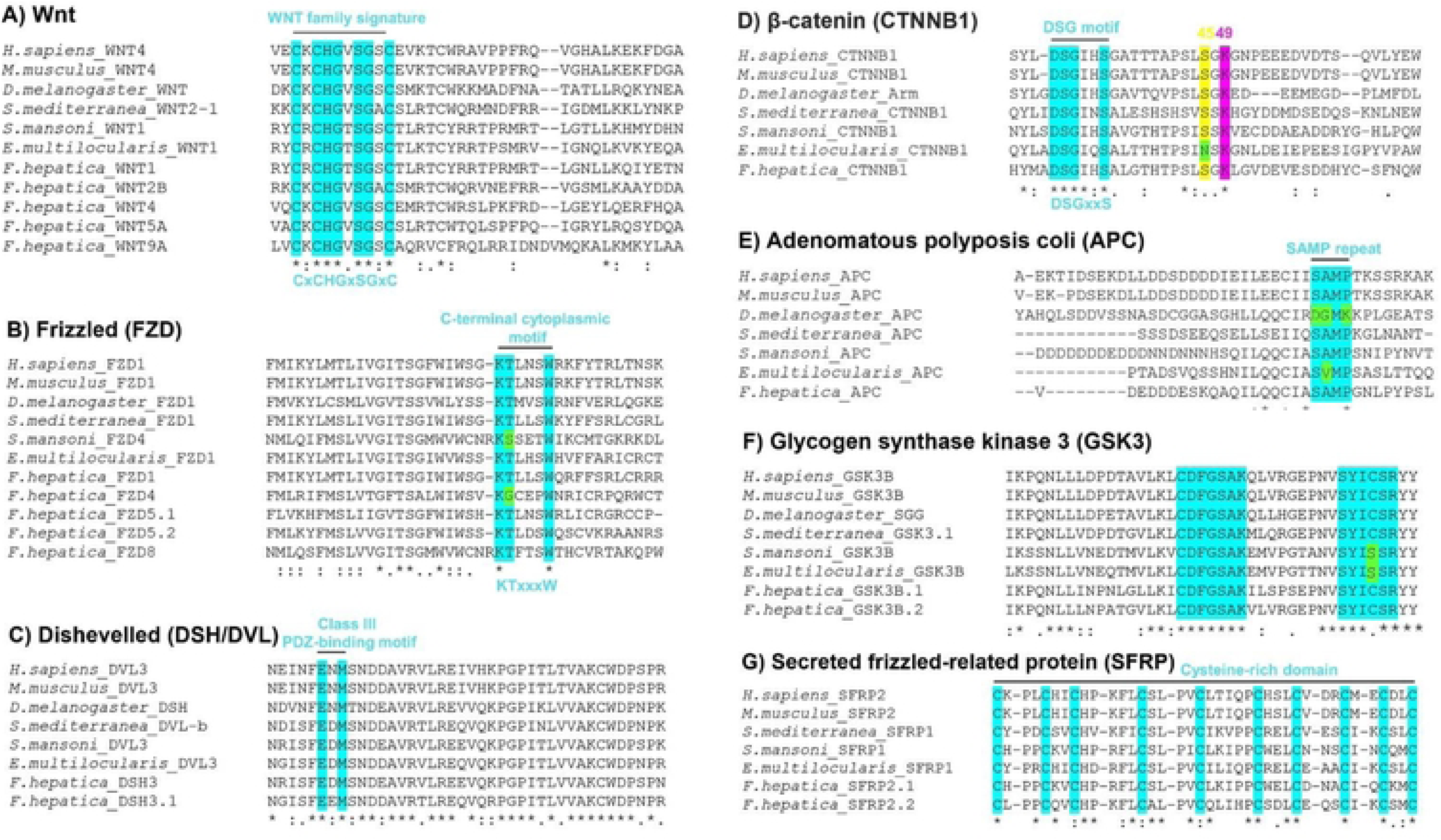
Liver fluke possess Wnt/p-catenin pathway components with conserved functional domains. Multiple sequence alignments demonstrate the conservation of key functional domains in putative Fasciola hepatica Wnt/p-catenin pathway components. A) Wnt family signature motif. B) C-terminal cytoplasmic motif of frizzled (FZD) receptors. C) Dishevelled class III PDZ-binding motif. D) p-catenin N-terminal residues essential for protein level regulation (light blue) and the conserved Ser45 (yellow) and Lys49 (magenta). DSG motif position denoted by black line. E) One of two adenomatous polyposis coli (APC) SAMP repeats. F) Two highly conserved motifs of the GSK-3 subfamily of serine/threonine protein kinases. G) SFRP cysteine rich domain. Residue positions relative to one another are not to scale. Sequence alignments were generated using Clustal Omega with default parameters. An asterisk (*) indicates a fully conserved amino acid, a colon (:) indicates conservation between strongly similar amino acids, a period (.) indicates conservation between weakly similar amino acids and a dash (-) indicates no consensus. Green denotes amino acid differences between the motifs of the species included.

#### Transcriptomic data support a role for Wnt/β-catenin signalling in juvenile *Fasciola hepatica* growth and development

Prior to the functional interrogation of individual pathway components, two distinct transcriptomic datasets were exploited in attempts to elucidate the relative importance of Wnt signalling in juvenile *F. hepatica,* the key life stage for control. Analyses of life stage transcriptome data [42] revealed that genes involved in active Wnt signalling were most highly transcribed in the early juvenile stages, notably 24 h old NEJs, while the expression of members of the Wnt/β-catenin destruction complex demonstrated bias towards adult stage fluke (Fig. 2) – both of these observations are consistent with elevated Wnt/β-catenin signalling in rapidly growing juveniles and reduced Wnt/β-catenin activity in adults. Few components involved in active Wnt/β-catenin signalling diverge from this trend, with the exception of dishevelled (DSH) which is most highly transcribed in the adult stage. The key effector of the pathway, β-catenin is also highly expressed in adult stage fluke, however, remains biased towards early juvenile stages.

**Figure 2.**
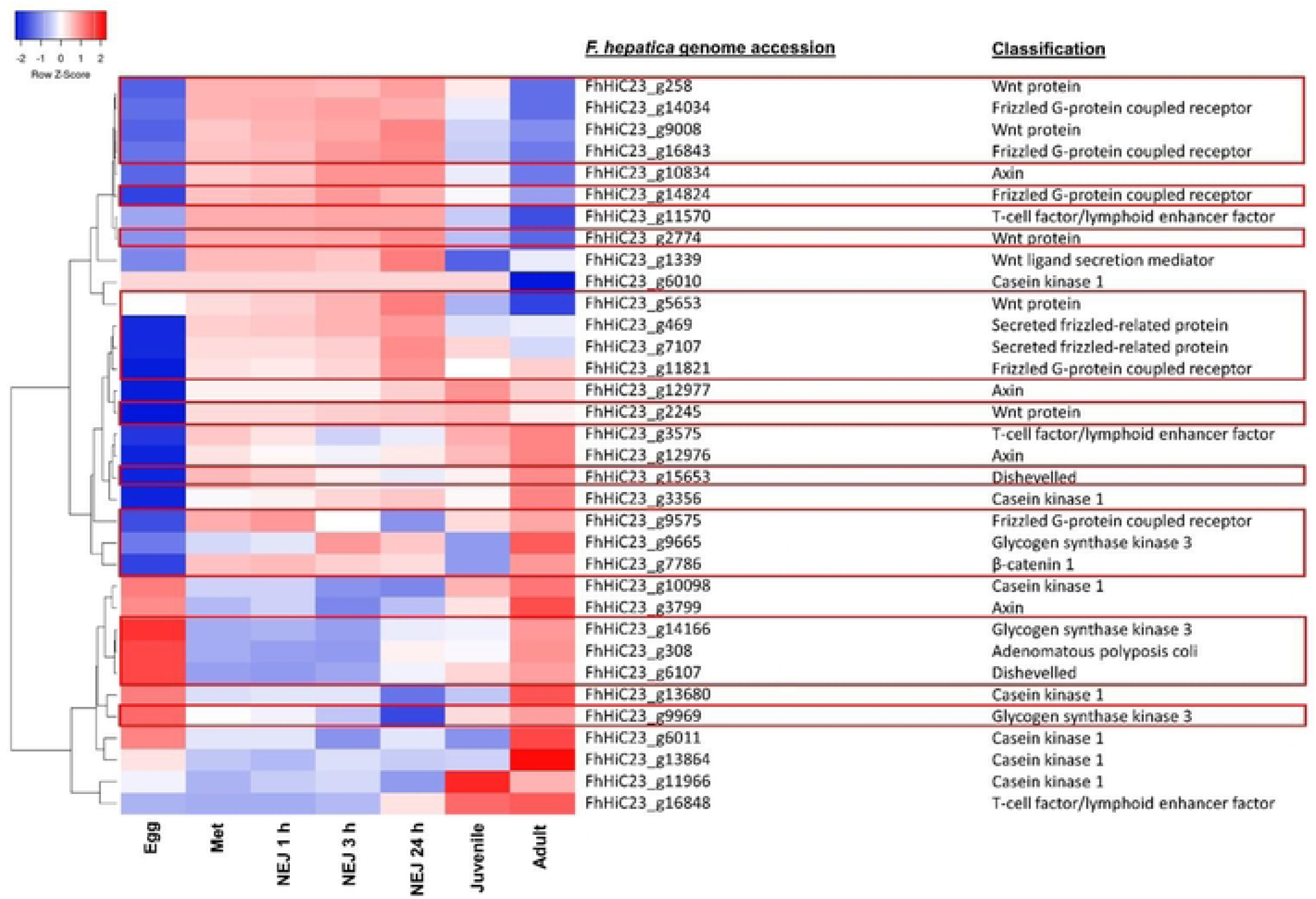
Wnt/p-catenin pathway components display marked changes in expression across the liver fluke life stages, with Wnt signalling being most active in juveniles. Developmentally staged expression heatmap generated from Iog2 TPM values of putative Wnt/p-catenin signalling pathway components. Columns correspond to life stages (egg; met - metacercariae; NEJ 1 h - newly-excysted juvenile 1 h post exeystment; NEJ 3h - NEJ 3h post-exeystment; NEJ 24 h - NEJ 24 h post-exeystment; Juvenile - 3-week-old worms collected from murine livers; Adult - adult worms collected from the bile ducts of bovine livers). Each row corresponds to a different pathway component, as denoted by gene ID and annotation. Red boxes denote genes targeted in RNAi experiments.

Previously, through comparative transcriptomic analyses, we demonstrated the downregulation of *Fh*Wnts and *Fh*FZDs in *ex vivo* three-week-old juveniles relative to developmentally delayed, time matched *in vitro*-maintained specimens [26], suggestive of key roles in early development. Following excystment, NEJs enter a period of remarkably rapid growth and development, as evidenced by a twofold increase in size every two weeks, with the concurrent upregulation of neoblast-like cell (cell-cycle) associated transcripts [48] and elaboration of the gut and reproductive structures [49]. As such, the upregulation of Wnt pathway components in these early developmental stages is likely reflective of the influential role of Wnt signalling in organogenesis and the formation of distinct cell and tissue types [50, 51, 52, 53, 54]. We would hypothesise that a key role for *Fh*Wnts during juvenile development is the establishment of anterior/posterior identity. While four *Fh*Wnts were differentially expressed in *in vitro* and *in vivo* juvenile fluke, just two *Fh*FZDs were downregulated in *in vivo* maintained juveniles [26]. This may represent promiscuity and interchangeability among *Fh*Wnt-FZD interactions, highlighting the potential for *Fh*Wnt signalling cascades to be highly complex and dynamic systems.

#### Localisation of *Fh*Wnt pathway components using fluorescence *in situ* hybridisation (FISH) reveals the partitioning of expression sites

This study localised eight *Fh*Wnt pathway component transcripts, revealing visually striking dichotomy among expression patterns throughout the *F. hepatica* body plan (Fig. 3). With the exception of the gut branches, *Fh*WNT1 appears to be absent from the posterior third of the worm and was predominantly detected within sub-tegumental parenchyma. *Fh*WNT2B transcripts were detected in a total of eight, relatively large cells comprising two distinct clusters in the anterior region of the worm. Despite their close proximity to one another, disparate morphologies indicate two unique cell types. Much like *Fh*WNT1, a strong signal also emanated from the guts of *Fh*WNT2B-labelled worms. While autofluorescence of the *F. hepatica* gut is a common occurrence following staining procedures, the guts of worms exposed to *Fh*WNT1 and *Fh*WNT2B sense probes were not highlighted, supporting the specificity of antisense probe binding and thus, the expression of *Fh*WNT1 and *Fh*WNT2B in the *F. hepatica* gut (S1 Fig). The expression loci of *Fh*WNT4 and *Fh*WNT5A transcripts both exhibit mid-anterior bias, being totally absent from the posterior region. *Fh*WNT4, localised to a network of cells scattered throughout the midbody region of the worm, while *Fh*WNT5A was detected within a population of cells across the mid-anterior region, in addition to parenchymal tissue surrounding the oral sucker. This curiously distinct band of cells/proportion of tissue highlighted by the *Fh*WNT5A antisense probe is perhaps reflective of a role in the patterning of the mediolateral axis, as is the case with a planarian WNT5 sub-family orthologue [15]. While the aforementioned *Fh*Wnt transcripts exhibit mid-anterior bias, *Fh*WNT9A transcripts localised to cells displaying a highly regimented pattern around the lateral edges of the posterior half of the worm. Widespread staining of the musculature was also detected in addition to a strong signal emanating from the parenchyma surrounding the oral sucker. While it is impossible to unequivocally identify and differentiate between specific cell types based on FISH alone, disparate morphologies and loci suggest that no two pathway components localised to the same cell population. As such, these data represent the most extensive and dichotomous Wnt expression patterns observed in platyhelminths to date. Leveraging the recently generated single cell RNAseq atlas of *S. mansoni* [55] revealed that homologues of all *Fh*Wnt pathway components localised using FISH were particularly enriched in various neuronal and muscle cell clusters, with sporadic expression in neoblasts, S1 progeny and flame cells. *Fh*CTNNB1 and *Fh*WNT1 homologues are ubiquitously expressed, corroborating the staining patterns observed in this study. In planaria, muscle cells are the source of developmental and positional control genes, including Wnts and SFRPs [56] and serve as landmarks during differentiation by relaying positional information to neoblasts. This, coupled with the *S. mansoni* homology data suggest that the majority of cell types expressing Wnt pathway transcripts in *F. hepatica* are muscle.

**Figure 3.**
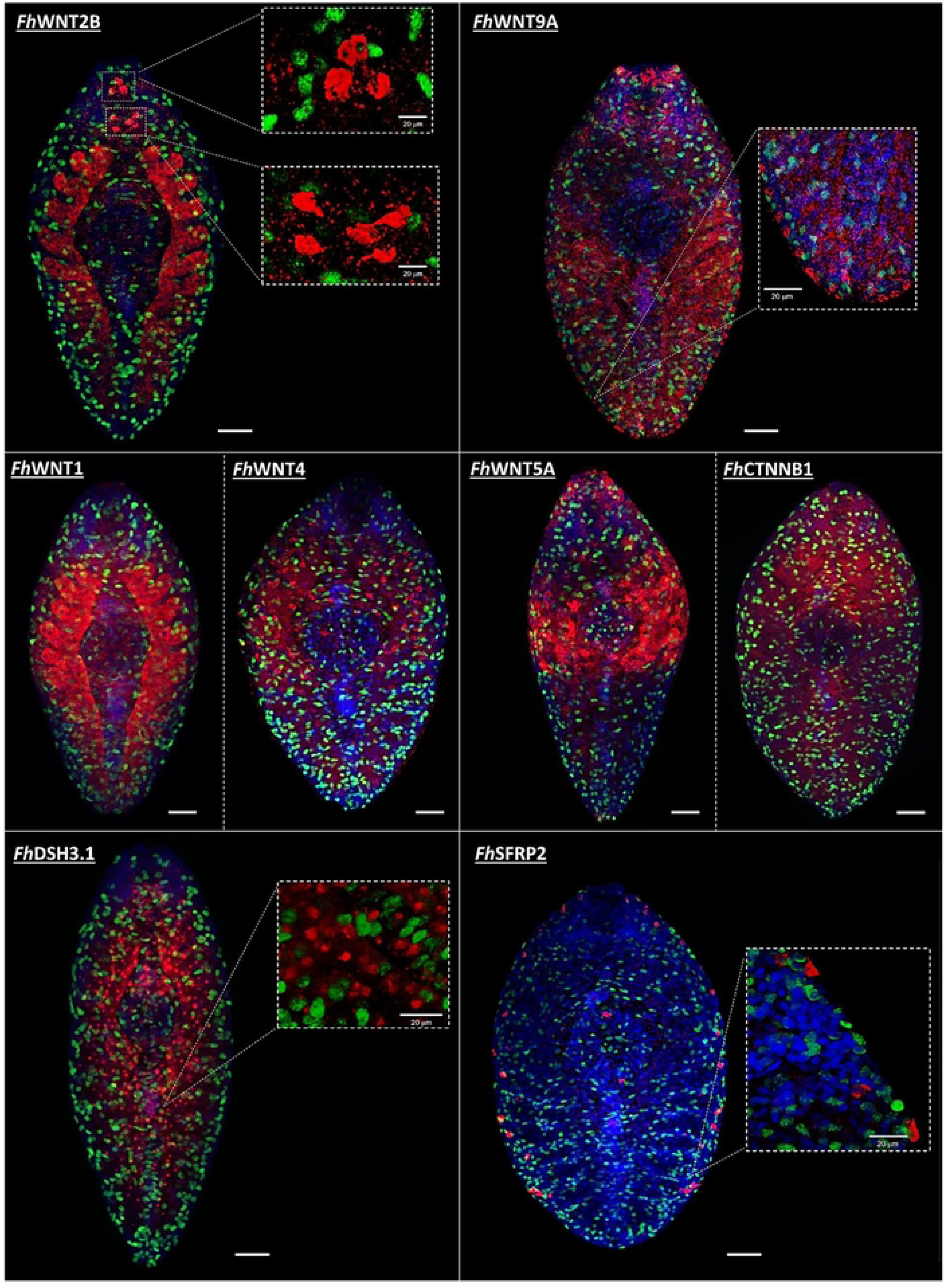
Juvenile liver fluke Wnt/p-catenin pathway components display distinct tissue expression patterns. Fluorescence *in situ* hybridisation (FISH)-mediated localisation of FfiWnt pathway transcripts in four-week old juvenile *Fasciola hepatica.* Target localisation is denoted by red (TAM RA) fluorescence, green fluorescence denotes EdU+ (neoblast-like) cells. DAPI (blue) served as a counterstain. Scale = 50 pm.

There exist multiple reports of Wnt pathway *in situ* hybridisation in planaria, the majority of which detail a posterior bias in Wnt protein ligand expression [15, 17, 52], while antagonists localise to the anterior pole [13, 15, 52]. The only *Smed*WNTs to diverge from this trend are a WNT2 subfamily member which is expressed laterally in the head and a WNT5 subfamily member demonstrating lateral expression along the dorsoventral axis [57, 58]. Wnt pathway transcript localisation in the *E. multilocularis* and *H. microstoma* metacestode stage mirrors this trend [22]. The highly dispersed spatial patterns of *Fh*Wnt transcript loci (Fig. 4) are, therefore, at odds with the strict anterior or posterior classification of Wnt antagonists and ligands reported in other flatworm species. *In vivo*, *F. hepatica* inhabit extremely caustic environments where the rapidity of growth and development is uncompromisingly vital and tissue damage commonplace. Therefore, it is plausible that Wnt signalling plays a more dynamic role in the biology of parasitic species than in the free-living members of the phylum. The dichotomy of expression patterns observed here may also be reflective of the involvement of Wnt signalling in the maintenance of stem cell niches [59, 60, 61].

**Figure 4.**
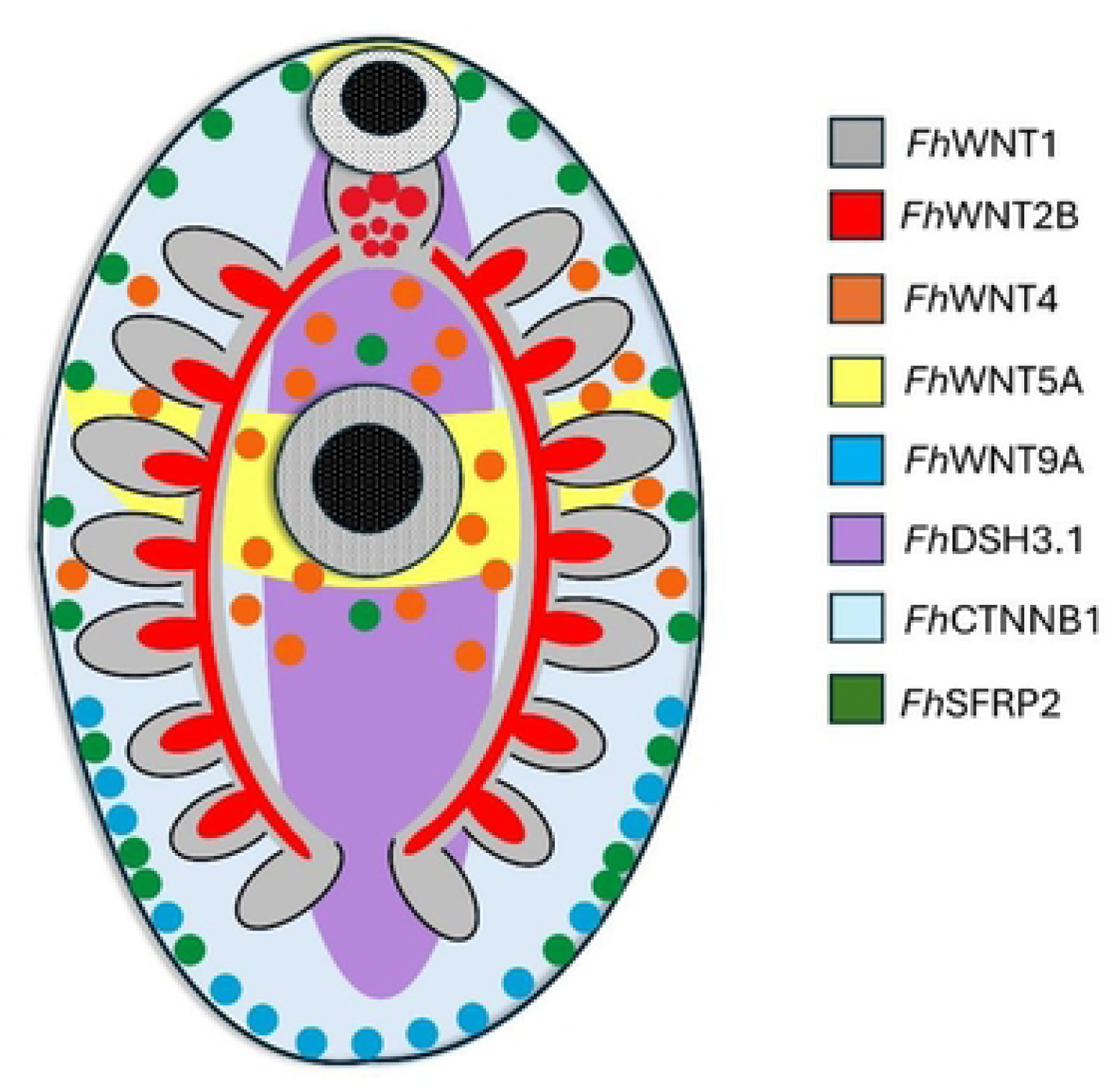
Schematic summarising the diverse expression sites of all Wnt/p-catenin pathway components successfully localised in juvenile Fasciol hepatica using fluorescence in situ hybridisation.

As anticipated due to its pleiotropic role, the expression of *Fh*CTNNB1 appears ubiquitous, localising throughout the parenchyma in a dorso-ventral manner. *Fh*DSH3.1 localisation revealed expression within a population of small cells, uniform in size along the anterior/posterior (A/P) axis of the worm and, accordingly, this transcript may be involved in the establishment of the A/P axis during development. The antagonistic *Fh*SFRP2 transcript localisation uncovered a remarkably symmetrical pattern, localising to discrete cells, likely muscle cell bodies, around the lateral edges of the worm in addition to a single cell above and below the ventral sucker. This is suggestive of a regulatory role in the establishment of bilateral symmetry during development. Although in many cases, Wnt targets localised in close proximity to EdU+ cells, the co-localisation of target Wnt pathway component transcripts and EdU within the same cell was not observed.

As their expression profiles are akin to *Fh*Wnts, the failure to localise *Fh*FZDs remains somewhat of an enigma. This discrepancy may be the result of FZD proteins being membrane-bound which, due to positional partitioning, may render their encoding transcripts less accessible to RNA probes than those targets within the extracellular matrix. Another possibility is the presence of secondary structures within *Fh*FZD mRNAs which may hinder RNA probe binding. However, as Wnt protein ligands exhibit limited diffusion in the extracellular environment and tend to act on neighbouring cells [43], it would seem logical to hypothesise that *Fh*FZD receptor loci and expression patterns would largely mirror those of their cognate *Fh*Wnt ligands.

#### RNA interference of *Fasciola hepatica* Wnt pathway components

In order to probe the role of Wnt signalling in juvenile *F. hepatica*, 18 canonical pathway components were selected for functional characterisation using RNAi-mediated gene silencing. Targets were prioritised based upon reports of differential expression in select transcriptomic datasets, plausibility as druggable targets and the existence of reported aberrant phenotypes in other organisms (see S3 Table for full list of RNAi targets and orthologues). All targets investigated were amenable to RNAi, as evidenced by significant target transcript knockdown relative to no dsRNA controls (S2 Fig.). It should be noted that due to consistent, significant die-off in *Fh*CTNNB1-silenced juveniles during the fourth week of RNAi trials, extractions were instead performed following 3 weeks of dsRNA exposures prior to the onset of mass death. Of the 18 targets silenced, 13 produced measurable growth phenotypes. With regards to *Fh*Wnts, knockdown led to significant reductions in worm growth for all five genes, evident from week one in the experimental setup (Fig. 5A, *n*=3; *Fh*WNT1, p=0.0253; *Fh*WNT2B, p=0.0359; *Fh*WNT4, p=<0.0001; *Fh*WNT5A, p=0.048; *Fh*WNT9A, p=0.0274). This inhibition of growth was sustained for the full duration of the trial, with *Fh*WNT4 yielding the most dramatic growth phenotype following four weeks of silencing (Fig. 5A, p=<0.0001). Interestingly, *Fh*WNT4 was also the most significantly differentially expressed WNT in the *in vitro* vs *in vivo* transcriptomes [26]. A conserved role fulfilled by the Wnt/β-catenin signalling cascade is the generation of various cell types, tissues and organ systems during development. In regenerating planaria, the silencing of Wnt protein ligands produces a plethora of aberrant phenotypes, including two-headed organisms, those void of tails, ectopic pharynges and the disappearance of both the anterior-posterior (A/P) and mediolateral axes [15, 57], illustrating functional diversity in platyhelminths. It is, therefore, unsurprising that the silencing of *Fh*Wnts profoundly inhibited the growth trajectories of juvenile *F. hepatica*.

**Figure 5.**
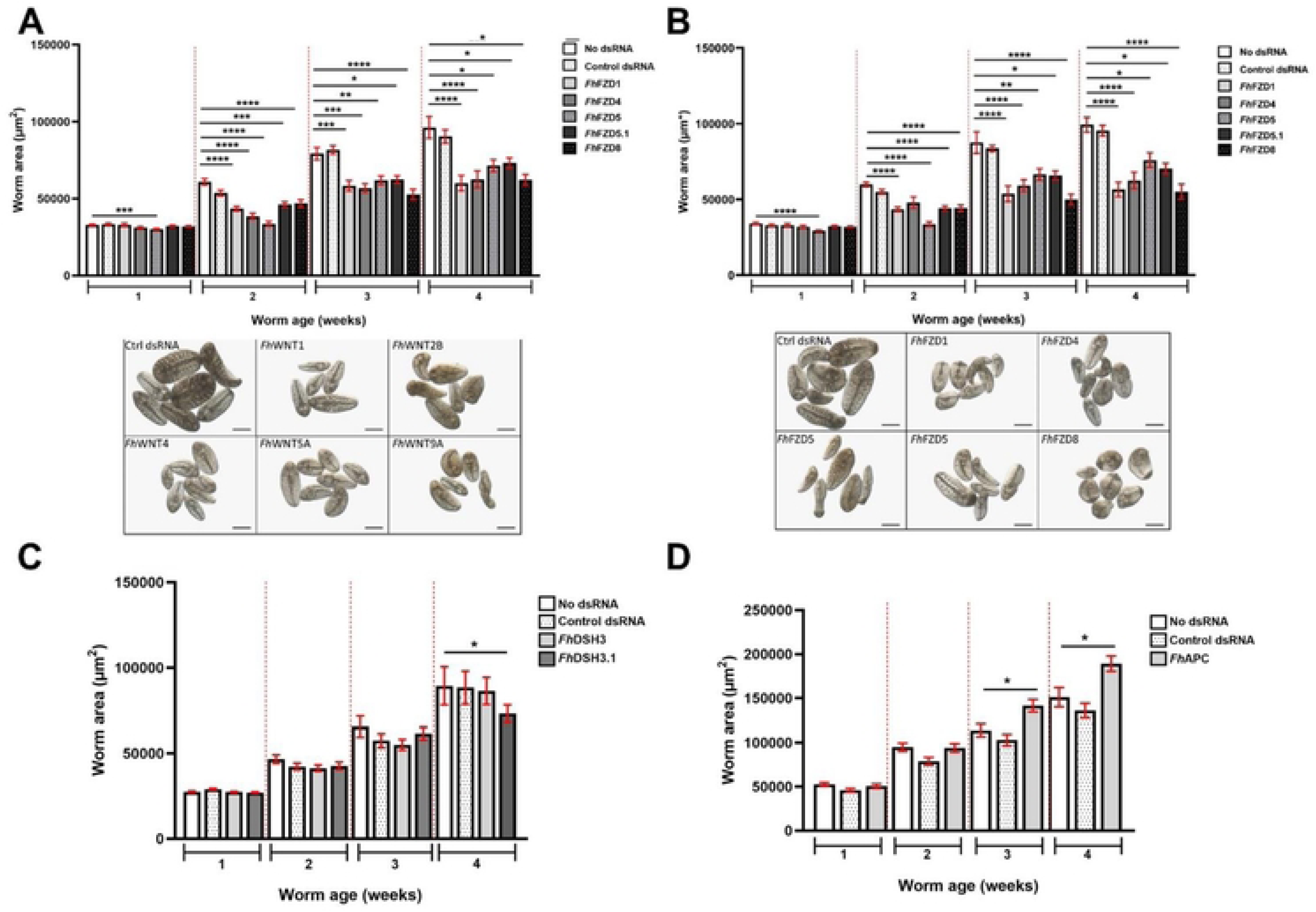
Silencing juvenile liver fluke Wnt/p-catenin pathway component gene transcripts dysregulates growth. Growth of juvenile Fasciola hepatica following four weeks of RNAi-mediated silencing of A) FhWnts B) FhFrizzleds in addition to light microscope images, C) FhDishevelled and D) FhAdenomatous polyposis coli. Worm area measured in pm^2^, data presented as pm^SEM (n=3). Scale = 100 pm. Statistical analyses were performed using Kruskal Wallis with Dunn’s post hoc tests. *, p<0.05; **, p<0.01; p<0.001; ****, p<0.0001.

Wnt protein ligands encourage the progression of myogenic progenitors throughout proliferative expansion and determine their terminal differentiation as both muscle cells and myogenic stem cells [62, 63]. They do so by inciting the transcriptional activation of myogenic regulatory factors including *Myf5* and *MyoD* [62] and as such, are heavily involved in myogenesis. *F. hepatica* possess three distinct types of muscle fibre (longitudinal, circular and diagonal) of which the major organ systems including the sub-tegumental musculature, gut, reproductive and attachment apparatus are comprised [64]. Indeed, the highly organised musculature constitutes the bulk of the worm’s mass. As such, the growth inhibition observed in *Fh*Wnt-silenced juveniles may principally be a consequence of dysregulated myogenesis. This is corroborated by the proposed localisation of *Fh*Wnts within various muscle cell types.

Wnt protein ligands also serve as niche factors [65], stimulating both the proliferation and self-renewal potential of tissue-specific stem cells [60]. Further, the silencing of β-catenin inhibits the proliferation of cancer stem cells by inducing cell cycle arrest [66, 67, 68]. This function appears to be conserved in *F. hepatica* with two *Fh*Wnts and *Fh*CTNNB1 exhibiting significant reductions in neoblast-like cell proliferation. As with *Fh*Wnts, a significant inhibition of growth was observed following the silencing of each *Fh*FZD. Unlike *Fh*Wnts, only one *Fh*FZD (*Fh*FZD5) exhibited a significant reduction in growth following one week of dsRNA exposures, however at week two, the significant inhibition of growth was observed in all *Fh*FZD targets (Fig. 5B). While the level of significance fluctuated in the cases of *Fh*FZD5 and *Fh*FZD5.1, reduced growth was still evident for all targets at week four of the experimental setup (*n*=3, *Fh*FZD1: p=<0.0001; *Fh*FZD4: p=<0.0001; *Fh*FZD5: p=0.0247; *Fh*FZD5.1: p=0.0463; *Fh*FZD8, p=<0.0008). As the activation of FZD GPCRs is contingent upon their association with a Wnt protein ligand, it is probable that the silencing of *Fh*FZDs phenocopied that of *Fh*Wnts and *vice versa*. The apparent absence of functional redundancy in *Fh*Wnts and *Fh*FZDs is an observation corroborated by the stark partitioning of *Fh*Wnt transcript expression observed in FISH.

Of two Dishevelled (DSH) orthologues, a reduced growth phenotype was only evident in *Fh*DSH3.1-RNAi worms (Fig. 5C). DSH regulates and channels Wnt signals into the various pathway branches and as such, the silencing of *Fh*DSH.3 likely inhibited growth via the disorganised funnelling of *Fh*Wnts into the canonical, or indeed, non-canonical branches. The silencing of DSH homologues in *S. mediterranea* leads to the loss of the A/P axis [58]. The localisation of *Fh*DSH3.1 transcripts along the anterior/posterior axis is also suggestive of a role in the maintenance of A/P identity in *F. hepatica*.

As aforementioned, neoblast-like cells drive growth and development in *F. hepatica.* Moreover, Wnt/β-catenin signalling is a key regulator of stem cell self-renewal, proliferation and differentiation. Taken alongside the observed reductions in growth exhibited by *Fh*Wnt/ β-catenin pathway component-silenced worms, it was hypothesised that neoblast-like cell activity would also be significantly reduced. To investigate this hypothesis, silenced juveniles were incubated in EdU for 24 h alongside those exposed to control dsRNA (Fig. 6A). This revealed that *Fh*WNT1- and *Fh*WNT2B-silenced juveniles displayed significantly reduced EdU+ (neoblast-like) cell numbers compared to control dsRNA-treated juveniles (Fig. 6D, p=0.0009 and p=0.0085, respectively). Of *Fh*FZD-silenced juveniles (Fig. 6B), only *Fh*FZD1 and *Fh*FZD4 caused significant reductions in neoblast-like cell proliferation (Fig. 6D, p=0.0478 and p=0.0152, respectively). As the results of this experiment mirrored those of *Fh*Wnts, in that two of five family members, when silenced, led to reduced cell proliferation, it may be hypothesised that *Fh*FZD1 and *FhFZD*4 constitute the receptors of *Fh*WNT1 and *Fh*WNT2B protein ligands. Despite silencing yielding growth phenotypes, neither *Fh*DSH3.1 or *Fh*APC affected neoblast-like cell proliferation. This appears somewhat surprising given that neoblast-like cells are proposed to drive growth and development in *F. hepatica* and related species [29, 49, 69]. In addition to its role in cell proliferation, Wnt signalling also mediates cell differentiation and fate determination, therefore, silencing the remaining *Fh*Wnts, *Fh*FZDs and *Fh*DSH3, may result in fewer cells of a specific cluster, manifesting as reduced growth in juvenile worms.

**Figure 6.**
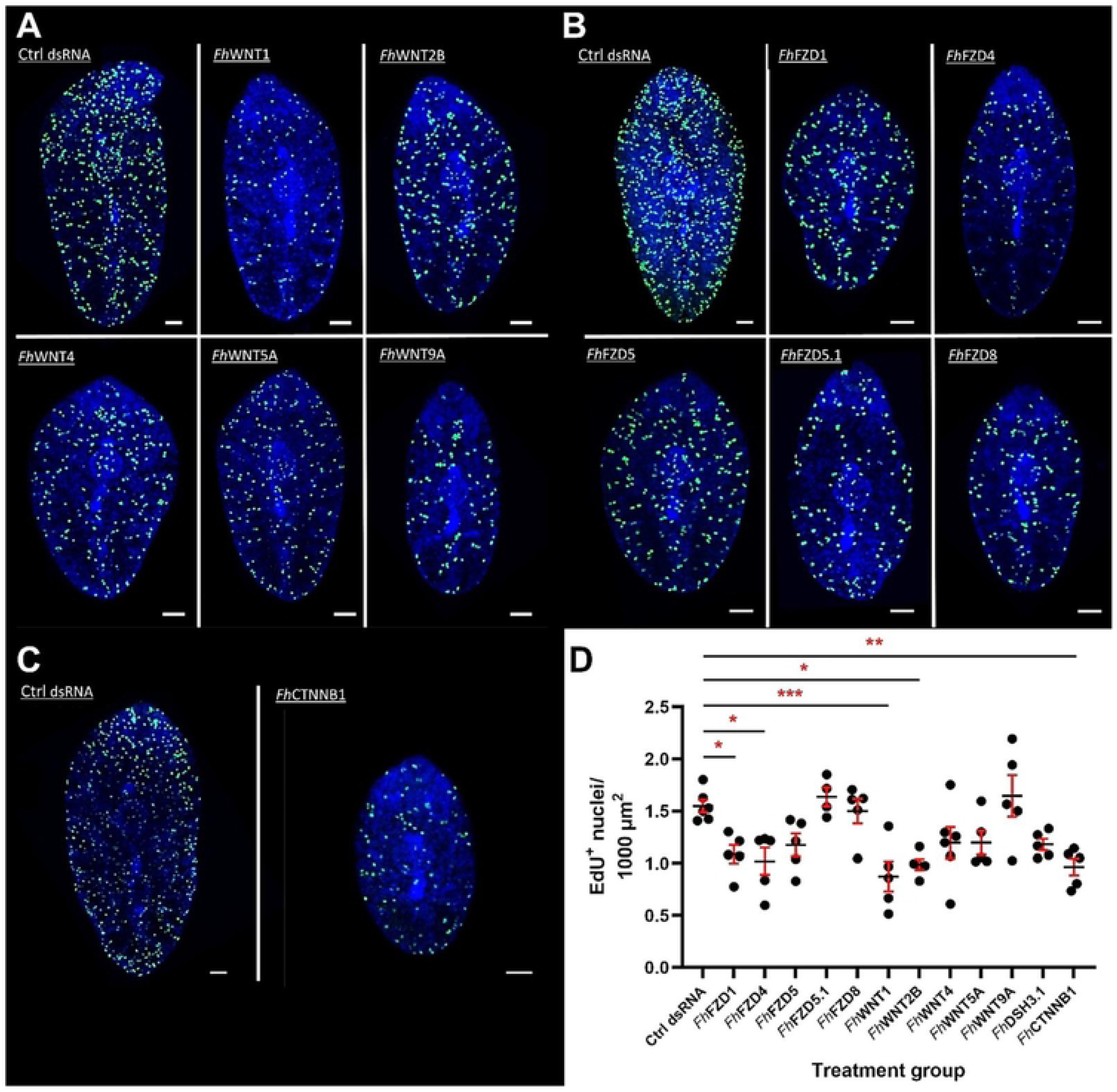
The silencing of selected juvenile liver fluke Wnt/p-catenin pathway component gene transcripts dysregulates neoblast-like stem cell proliferation. Maximally projected confocal z-stack images of EdU+ nuclei (green) in juvenile *F hepatica* following four weeks of silencing A) F/?Wnts, B) FfiFrizzleds and C) F/?CTNNB1. DAPI (blue) served as a counterstain. Scale = 50 pm. D) Neoblast-like cell proliferation in FfiWnt pathway component silenced juveniles yielding reduced growth phenotypes relative to dsRNA control groups. Data presented as mean EdU+ nuclei ± SEM (n=3). Statistical analyses were performed using a one-way ANOVA with Dunnett’s post hoc tests. *, p<0.05; **, p<0.01; ***, p<0.001.

As Wnt/β-catenin signalling is negatively regulated by the presence of a destruction complex, it was hypothesised that silencing components of this complex would remove regulatory pressure, thereby leading to increased growth. Despite significant levels of transcript knockdown, no growth phenotypes were observed in any of the destruction complex targets, with the exception of *Fh*APC (S3 Fig, Fig. 5D). At week four, *Fh*APC silenced juveniles were significantly larger than control groups (Fig 5D, *n*=3, p<0.05). The most common mutation preceding the development of colon cancer is the inactivation of APC which, under normal conditions serves as a tumour suppressor via the modulation of cytoplasmic β-catenin levels. This, coupled with the fact that APC orchestrates the Wnt/β-catenin destruction complex assembly and function, may explain why it was the sole destruction complex component yielding a phenotype. The importance of APC has also been demonstrated in regenerating planaria where silencing results in the formation of a tail in place of a head [13]. The lack of phenotypes arising from *Fh*GSK3B-silencing may be explained relatively simply in that the generation of life stage expression profiles for *Fh*Wnt/β-catenin pathway components revealed that members of the destruction complex exhibit their lowest levels of expression in early juvenile stage fluke. As the RNAi screen was performed on fluke spanning those very age groups, knocking down the transcripts of *Fh*GSK3Bs further had no discernible effect.

Following FISH, a juxtaposition was identified between the expression patterns of *Fh*SFRP2 and *Fh*WNT9A, perhaps suggestive of a functional relationship. While neither *Fh*SFRP agonists yielded growth phenotypes, *Fh*SFRP2-silenced juveniles exhibited a significant increase in motility (S3 Fig., *n*=3, 199.1±22.65%, p=<0.0001). However, it has been suggested that SFRPs may not serve as Wnt antagonists in platyhelminths due to the absence of a netrin-like domain required for their anchorage to the extracellular matrix [70]. As such, the observed increase in motility may be the result of an unknown function executed by *Fh*SFRP2 in *F. hepatica*.

#### *Fh*CTNNB1 is essential for the development of the juvenile *Fasciola hepatica* neuromuscular system

The phenotypic consequences of silencing *Fh*CTNNB1 were undoubtedly the most profound observed in this study. This is supported by the ubiquitous expression of the *Fh*CTNNB1 transcript, an observation also reported in planaria [71, 72]. Significant reductions in worm growth were evident in two-week-old juveniles, an effect that was further exaggerated at weeks three and four (Fig 7A, *n*=3, p=0.0009, p=0.0009, p=0.0003, respectively), accompanied by concomitant reductions in neoblast-like cell proliferation (Fig. 6C, p=0.0020). Interestingly, the silencing of *Fh*CTNNB1 also resulted in significant fluke die off, with average survival being just 19% at week four (Fig. 7B, *n*=3, 19±5.85%, p=<0.0001). Worm death was defined by a darkened appearance, gut oedema and total loss of tissue integrity within 24 h (Fig. 7C). As the key effector of the canonical signalling pathway, β-catenin regulates gene transcription and triggers conformational changes in chromatin [73, 74], thereby acting as a master regulator, fulfilling a myriad of context-dependent roles during development, regeneration and homeostasis. In vertebrates, the most prominent role of β-catenin during development is in the anteroposterior patterning of the central nervous system [75].

**Figure 7.**
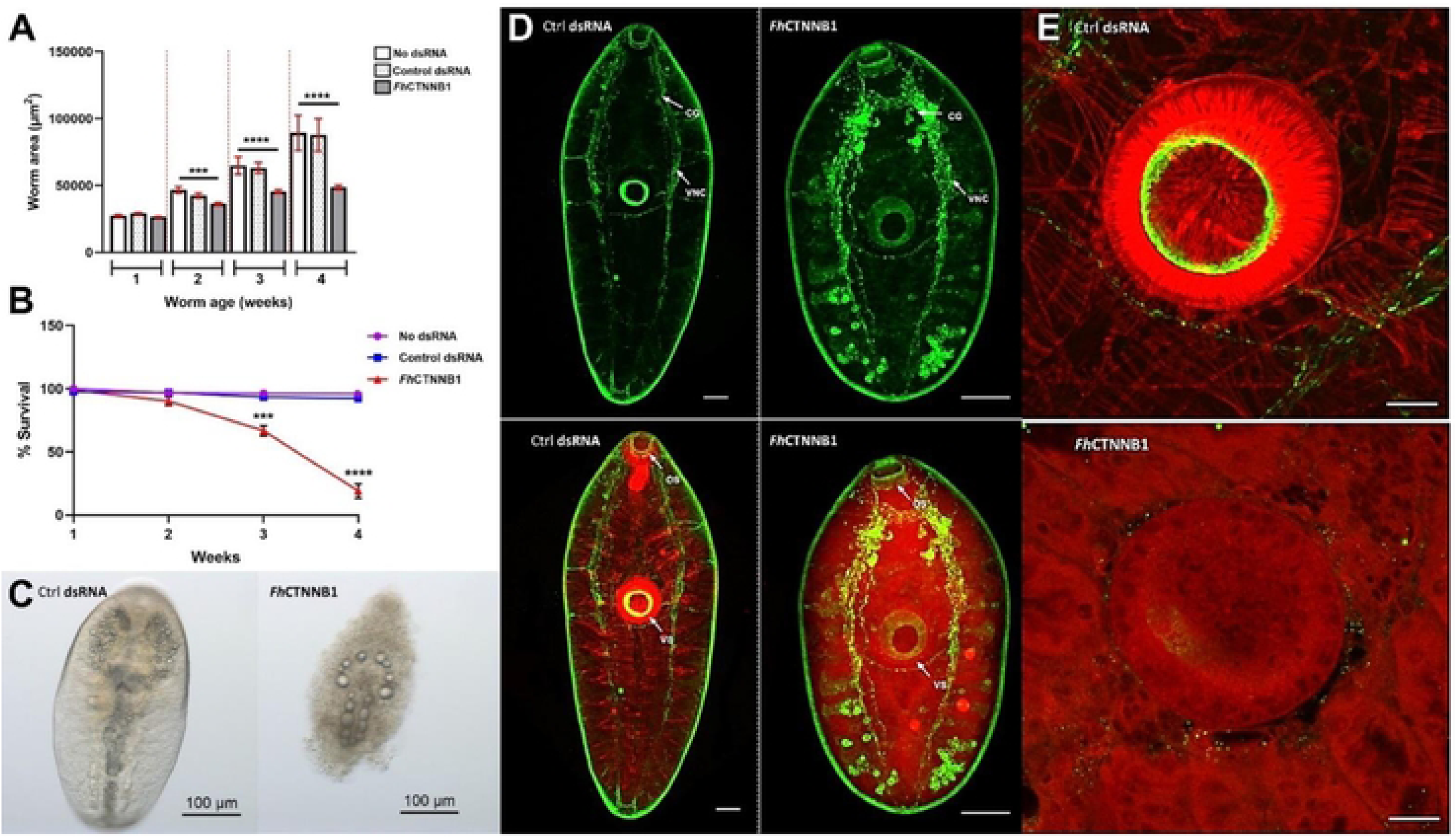
*Fh*CTNNB1 silencing in juvenile liver fluke dysregulates muscle/nervous system structure/development, inhibits growth and reduces survival A) Growth following four weeks of RNAi-mediated F/vCTNNBI-silencing. Worm area measured in pm^2^, data presented as pm^SEM (n=3). B) Survival. Data show mean survival (±SEM) relative to untreated control (RPMI) juveniles. Statistical analyses were performed using unpaired t-tests and Kruskal Wallis with Dunn’s post hoc tests. ***, p<0.001; ****, p<0.0001. C) Light microscope images demonstrating the morphological features of *Fh*CTNNB1-silencing induced death in a three-week old F/?CTNNB1-silenced juvenile compared to a time-matched control dsRNA-treated juvenile. D) Confocal scanning laser micrographs of wholemount dsRNA control and *Fh*CTNNB1 silenced three-week old *F hepatica* subjected to ICC. Green staining denotes NPF immunoreactivity, highlighting the nervous system, while TRITC-labelled phalloidin (red) highlights filamentous-actin. CG, cerebral ganglia; VNC, ventral nerve cord; OS, oral sucker; VS, ventral sucker. E) Phalloidin-labelling of the filamentous-actin structure of the musculature comprising the ventral suckers (red) of dsRNA control and *Fh*CTNNB1 silenced three-week old *Fasciola hepatica.* Scale = 50 pm.

To investigate the phenotypic consequences of *Fh*CTNNB1 knockdown in greater detail, silenced and control specimens were subjected to neuropeptide F (NPF) immunocytochemistry (ICC) and fluorophore-phalloidin staining in order to visualise a significant component of the peptidergic nervous system and musculature arrangements, respectively. *Fh*CTNNB1-silenced worms were prepared for ICC at three weeks old prior to the onset of mass die off. Indeed, *Fh*CTNNB1-silenced juveniles exhibited aberrant, uncoordinated development of the central nervous system, with unusually dense NPF-immunoreactivity in both the cerebral ganglia and ventral nerve cords, and the formation of spherical, ectopic neuronal cell bodies (Fig. 7D). Such observations could be deemed consistent with hypercephalisation, a phenomenon also reported in *S. mediterranea*, where the silencing of CTNNB1 produces organisms exhibiting ectopic eyes, radial-like hypercephalisation and a loss of A/P identity [71].

Phalloidin staining of F-actin revealed an equally striking visual phenotype, demonstrating a loss of filamentous-actin patterning and structural integrity within the musculature of *Fh*CTNNB1 silenced juveniles (Fig. 7D). This was particularly evident when examining the highly muscular oral and ventral suckers (Fig. 7E). In addition to its role in signal transduction, β-catenin also serves as a scaffold molecule and is a key constituent of cadherin-based adherens junctions [76]. Adherens junctions are unique macromolecular structures, vital in both intercellular communication and adhesion [77]. Their adhesive competence is dependent upon members of the cadherin superfamily which bind homotypically to complementary cadherin molecules on neighbouring cells [78]. The direct interaction of β-catenin with the cytoplasmic domain of cadherins prompts the establishment of a complex with α-catenin and the subsequent linkage of actin filaments to adherens junctions, forming a structural continuum [79]. In this respect, these specialised junctions assist in the formation of the polarised epithelial tissues required for morphogenesis and organism integrity [80]. The significance of this architectural role was elegantly illustrated by the inability of β-catenin-deficient murine embryonic stem cells to differentiate into a mesodermal germ layer and form a neuroepithelium as a result of flawed cell-cell adhesion [81]. Assuming β-catenin performs a dual role in *F. hepatica*, the silencing of *Fh*CTNNB1 may preclude the formation of cellular junctions, leading to an imbalance in the structural properties of the cytoskeleton and thereby, loss of tissue integrity and death. The proposed bifunctionality may also explain why *Fh*CTNNB1 was not differentially expressed in *ex vivo* juveniles, despite the downregulation of *Fh*Wnts and *Fh*FZDs [26].

While this study presents both transcriptomic and phenotypic evidence lending support to the hypothesis that Wnt signalling is a key molecular driver of growth and development in juvenile *F. hepatica*, similar studies in related species oppose such findings. A large-scale RNAi screen performed in *S. mansoni* silenced four FZD orthologues (Smp_118970, Smp_139180, Smp_155340, Smp_174350), in addition to a β-catenin-like protein (Smp_089400) with no measurable phenotypes following 30 days of intermittent dsRNA exposures [81]. This appears somewhat peculiar given the close phylogenetic relationship between *F. hepatica* and *S. mansoni*. Quite simply, this apparent disparity may be a consequence of the RNAi screen of Wang *et al* [82] employing fully developed, adult parasites. Further, target expression analyses were not conducted, therefore, the absence of phenotypic readouts could be due to the absence of, or modest transcript and/or protein knockdown. As schistosomulae are amenable to RNAi [83, 84], silencing Wnt pathway components in this earlier life stage may yield tangible phenotypes.

#### Pharmacological inhibition of *Fh*Wnt/β-catenin pathway components

Due to the identification of robust links between aberrant Wnt signalling and tumorigenesis, recent years have observed a concerted effort within the field of oncology to exploit small-molecule inhibitors targeting various Wnt pathway components as novel anti-cancer treatments. Pyrvinium pamoate (PP) is an FDA-approved anthelmintic, originally employed in the treatment of *Enterobius vermicularis* (pinworms). More recently, PP was found to promote the degradation of β-catenin by selectively potentiating casein kinase 1α activity in various cancer cell lines, inhibiting tumour growth via the downregulation of Wnt signalling [85, 86]. Similarly, biweekly exposures to PP significantly impeded growth and neoblast-like cell proliferation in juvenile *F. hepatica* (Fig. 8B & 8C, *n*=3, p<0.0001, Fig. 8E, *n*=3, p=0.0114, respectively). After just one week, the degree of growth inhibition observed in PP-treated juveniles paralleled that of TCBZ. Moreover, concentrations greater than 0.1 µM resulted in a loss of tissue integrity and proved lethal. Due to significant die off in 0.1 µM PP-treated juveniles during week three, the experiment had to be terminated. PP-induced fatalities have also been reported in *S. mansoni* schistosomula [87].

**Figure 8.**
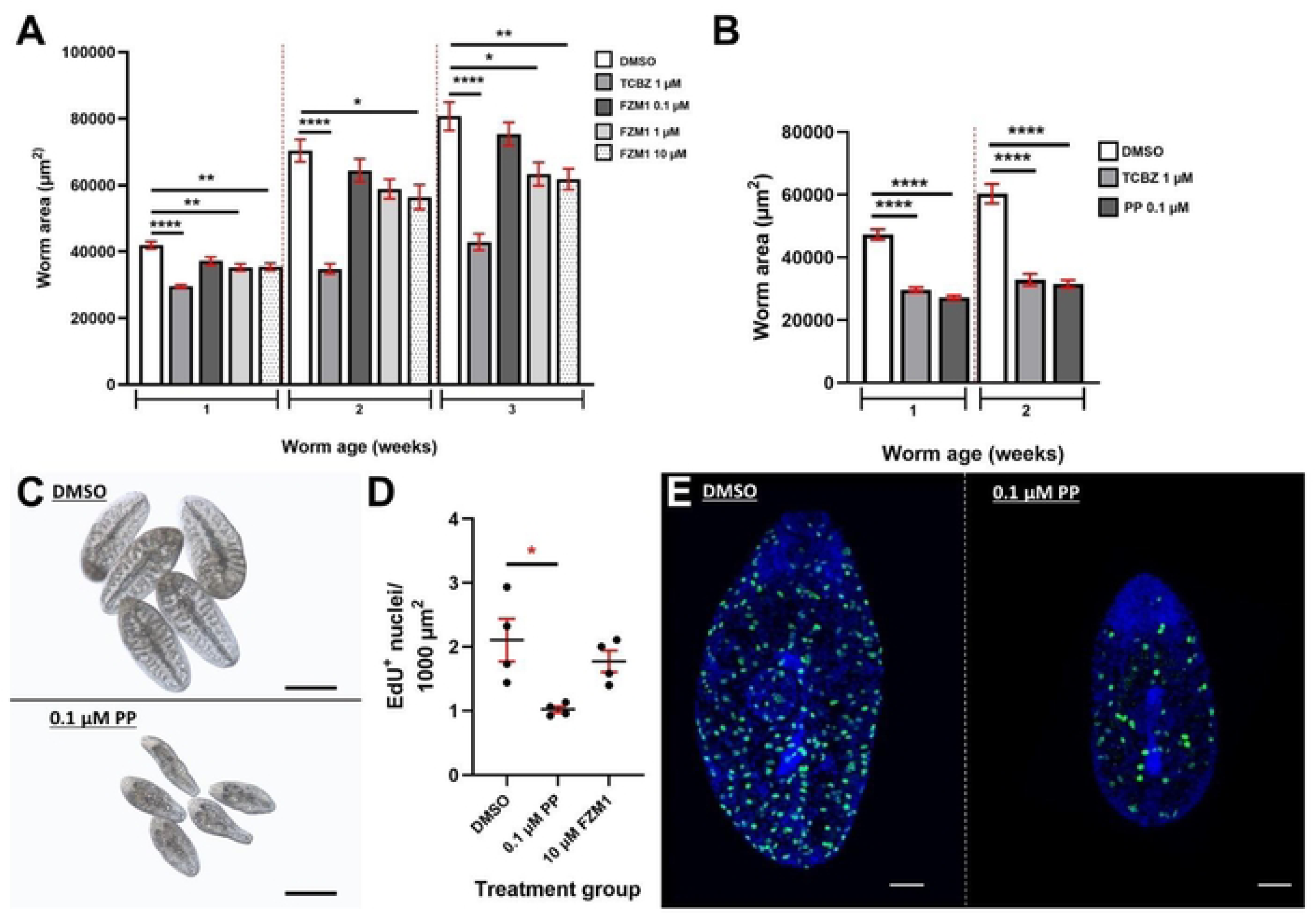
Selected Wnt/p-catenin pathway antagonists dysregulate neoblast-like cell proliferation and growth in juvenile liver fluke. A) Growth of juvenile *F hepatica* following biweekly exposures to A) FZM1 and B) Pyrvinium pamoate (PP). TCBZ served as a positive control. Worm area measured in pm^2^, data presented as pm^SEM (n=3). Statistical analyses were performed using Ordinary One-way ANOVA or Kruskal Wallis with Dunn’s post hoc tests. *, p<0.05; p<0.01; ****, p<0.0001. C) Light microscope images showing size and aberrant morphology of two-week old 0.1 pM PP-treated juveniles relative to DMSO controls. D) Neoblast-like cell proliferation in juvenile *Fasciola hepatica* following treatment with PP and FZM1 relative to DMSO controls. Data presented as mean EdU+ nuclei ± SEM. Statistical analyses were performed using a one-way ANOVA with Dunnett’s post hoc tests. *, p<0.05. C) Maximally projected confocal z-stack images of EdU+ nuclei (green) in DMSO control and PP-treated two-week old juvenile *F hepatica.* DAPI served as a counterstain (blue). Scale = 50 pm.

The inhibition of Wnt signalling is known to interfere with the recruitment of stem cells to wound sites following mechanical injury [88, 89] and disrupts their proliferative potential during the healing process [90]. Moreover, it has been demonstrated that TCBZ enters *F. hepatica* tissues via trans-tegumental diffusion [91], with tegumental damage proving fatal [92]. If PP does indeed inhibit Wnt signalling in *F. hepatica*, the dysregulation of neoblast-like cells coupled with compromised musculature would likely result in the accumulation of ultrastructural damage to the tegument. This in turn, would increase the rapidity and ease of TCBZ diffusion into tissues, enhancing the systemic effects of the drug. Further, reductions in the proliferative potential of neoblast-like cells would render worms less equipped to defend themselves from drug action.

Although not unequivocally determined, PP is postulated to inhibit glucose uptake and mitochondrial fumarate reductase in pinworms [93]. Since Wnt activation has been shown to modulate mitochondrial function which, in a feedforward loop, helps to drive Wnt signalling in human cells [94], the possible dysregulation of multiple pathways and processes contributing to the observed PP-induced phenotype in *F. hepatica* cannot be ruled out. Biweekly exposures to FZM1, a FZD4 inhibitor also significantly reduced juvenile fluke growth (Fig. 8A, *n*=3, p=0.0066), again, phenocopying the effects of RNAi-mediated silencing of *Fh*FZD4. FZM1 binds to an allosteric binding site within intracellular loop 3 of FZD4, resulting in conformational changes within the receptor which ultimately inhibit Wnt/β-catenin signalling [95].

## CONCLUSION

In the face of widespread flukicide resistance, liver fluke control is precarious, such that the discovery and validation of novel drug targets is imperative. This study represents the first functional characterisation of a developmental signalling pathway in *F. hepatica*, highlighting components of the Wnt/β-catenin (canonical) signal cascade as key molecular drivers of growth, development and cell proliferation, biological processes essential for parasite propagation and survival *in vivo*. Notably, in addition to reductions in growth and stem cell proliferation, the silencing of *Fh*CTNNB1 led to aberrant development of the neuromuscular system, a well-established target of many existing anthelmintics. FISH-mediated localisation of *Fh*Wnt pathway component transcripts revealed dichotomous expression patterns and corroborate the results of reverse genetics experiments. We have demonstrated the efficacy of Wnt pathway inhibitors against juvenile *F. hepatica in vitro*, with the effects of PP and FZM1 phenocopying RNAi results. This provides impetus for larger scale compound screens targeting conserved developmental pathways with the view of re-purposing these drugs as novel flukicides. Further, combination therapies employing small molecule inhibitors in conjunction with existing flukicides, may synergise drug effects, thereby offering respite in field cases of TCBZ resistance. This is an avenue that warrants further investigation. Collectively, these data have advanced our understanding of the molecular mechanisms underpinning pathogenic juvenile *F. hepatica* growth, development and neoblast-like cell dynamics. Dysregulating Wnt/β-catenin signalling in juvenile fluke would undermine parasite virulence such that its pathway components represent attractive targets for the development of novel flukicides.

## MATERIALS AND METHODS

### *In silico* characterisation of *Fasciola hepatica* Wnt signalling pathways

Putative homologues of all major components of both canonical and non-canonical Wnt signalling pathways were identified in *F. hepatica* using a reciprocal BLAST (basic local alignment search tool) approach. Sequences from *Homo sapiens*, *Mus musculus*, *Drosophila melanogaster* and *Caenorhabditis* elegans were obtained from Uniprot for each gene of interest (see S1 File for query sequence gene accessions) and queried against the *F. hepatica* genomes (PRJEB25283/PRJEB58756 and PRJNA179522) available on WormBase ParaSite v16 using BLASTp and tBLASTn with default parameters. Returned hits (sorted by E-value) were subsequently screened against the NCBI non-redundant (nr) protein sequence database using BLASTp to validate sequence identity. The organism field was set to exclude Platyhelminthes. Where the top hit was not the gene of interest, that sequence was omitted from the dataset.

As an additional measure of confidence, putative *F. hepatica* Wnt, frizzled (FZD), β-catenin, dishevelled (DSH) and glycogen synthase kinase 3 (GSK3) protein sequences were aligned with those of *H. sapiens*, *M. musculus*, *D. melanogaster*, *S. mediterranea* and *S. mansoni* orthologues in Clustal Omega using ClustalW with character counts. The integrity and classification of homologues was further examined through the identification of conserved domains with the aid of Pfam HMMScan (https://www.ebi.ac.uk/Tools/hmmer/search/hmmscan) and Interpro (www.ebi.ac.uk/interpro/search/sequence-search). The presence of functional domains and characteristic motifs were considered to be evidence of gene conservation and functional viability.

To gain insight into the degree of expression of various pathway components and thus, the relative importance of Wnt signalling throughout the *F. hepatica* life cycle, the life-stage transcriptomic data supporting the *F. hepatica* genome project was mined (available on WormBase Parasite v16; egg (*n*=1), metacercariae (*n*=3), 1-hour newly excysted juvenile (NEJ, *n*=2), 3-hour NEJ (*n*=2), 24-hour NEJ (*n*=2), 21-day old juvenile (*n*=1) and adult-stage (*n*=1) *F. hepatica*) to obtain median transcripts per million-mapped reads (TPM) values for all *F. hepatica* canonical and non-canonical Wnt pathway components. TPM values were subsequently transformed to a Log2 scale and uploaded to heatmapper (http://www.heatmapper.ca/) [96] set for Average Linkage, and Pearson Distance Measurement to generate a life stage expression heatmap.

### Excystment and *in vitro* maintenance of juvenile *Fasciola hepatica*

Experiments were carried out using the Italian (TCBZ-susceptible) *F. hepatica* isolate (Ridgeway Research Ltd, UK). Encysted metacercariae were excysted as described by McVeigh *et al.* [97]. Our excystment protocol is available, in full at https://www.protocols.io/private/B165F2D0C64B11EDAB840A58A9FEAC02. In short, encased metacercariae were popped from their outer cyst walls and treated with 10 % sodium hypochlorite for 2-3 min prior to the addition of excystment solution. Metacercariae were incubated in excystment solution at 37°C for one hour. Newly excysted juveniles (NEJs) were washed in warmed RPMI 1640 (#11835105, ThermoFisher Scientific), before being transferred to either 40 µl of dsRNA (if destined for RNAi experiments) or 200 µl of 50 % chicken serum (#16110082, Thermo Fisher Scientific) in RPMI (v/v) (CS50) in round bottomed, 96 well plates (Sarstedt). Juvenile fluke were maintained for the duration of experiments in groups of ∼20 worms per well in a humidified incubator set to 37°C with 5% CO2 atmosphere. Culture medium was replaced every three days.

### Whole mount fluorescence *in situ* hybridisation (FISH)

Juvenile worms were excysted as above and maintained under standard culture conditions for four weeks. On day 27 of culture, worms were incubated in 500 µM 5-ethynyl-2′-deoxyuridine (EdU) in CS50 for 24 h to enable the co-localisation of FISH targets with EdU^+^ (neoblast-like) cells. At four weeks, worms were washed in RPMI and anaesthetised via a three-minute incubation in 0.25 % tricaine (ethyl 3-aminobenzoate methanesulfonate, #A5040, Sigma-Aldrich). From this point, RNase-free pipette tips were used, and reagents were made using diethyl pyrocarbonate (DEPC)-treated RNase-free water (this is applicable to all steps until post Riboprobe incubation). Worms were transferred to a 20 µl droplet of 4% formaldehyde (36% methanol formaldehyde [Sigma Aldrich], diluted in PBSTx (1x phosphate buffered saline, PBS) with 0.3% Triton^TM^ X-100 [Sigma-Aldrich]) in a petri dish and flat fixed under a weighted coverslip for 10 min. Following removal of the coverslip, worms were collected into NoStick® hydrophobic microtubes (#1210-S0, SSIbio) containing 1 ml 4% formaldehyde, and free fixed for a further 10 min. After fixation, worms were washed in PBSTx before being dehydrated via a 10-min incubation in 1:1 PBSTx:Methanol (MeOH). Worms were stored in 100% MeOH at −20°C until use. All incubation and wash steps were carried out under rotation at room temperature (RT).

The synthesis of DIG-labelled RNA probes was achieved through *in vitro* transcription reactions. Amplicons labelled with the T7 promoter sequence; 5’-TAATACGACTCACTAT AGGGT-3’ were generated with end-point PCRs using FastStart Taq Polymerase (#4738357001, Sigma Aldrich) and purified using the ChargeSwitch™ PCR Clean-Up Kit (#CS12000, ThermoFisher Scientific). Primer sequences can be found in S1 table. Transcription reactions consisted of 2 µl transcription buffer, 1 µl T7 polymerase (T7 RNA polymerase kit, ThermoFisher Scientific), 1 µl digoxigenin-labelled ribonucleotide mix (Roche) and 6 µl DNA template. Probes were then incubated overnight at 27°C before being DNase treated for 15 min at 37°C (0.25 µl DNase I, 0.25 µl Transcription Buffer and 2 µl RNase-free H2O). Probes were precipitated overnight at −80°C with the addition of 1.25 µl 4 M LiCl and 37.6 µl chilled molecular grade ethanol [98]. Post-precipitation, RNA was pelleted via a 20-min centrifugation (x16,000 *g* at 4°C). The pellet was washed in 70% chilled molecular grade ethanol and centrifuged for a further 5 min before discarding the supernatant and allowing to air dry for 15 min. The pellet was then resuspended in 20 µl RNase-free H2O and probe concentrations and purities were measured using a DeNovix DS-11 FX spectrophotometer/fluorometer. Concentrations were adjusted to 50 ng/µl via the addition of hybridisation solution (25 ml de-ionized formamide, 12.5 ml 20X SSC, 100 µl of 50 mg/ml yeast RNA, 5 ml 10% (v/v) Tween-20, 5 ml 50% Dextran Sulfate [Sigma Aldrich] and 2.4 ml DEPC-treated H2O) and probes were stored at −20°C until use. Yeast RNA was generated as described by Jing [99].

The core FISH protocol employed here (including the preparation of all stock solutions) was that detailed by King & Newmark [98]. The only modifications made were in the treatment of parasites pre-hybridisation and are as follows; worms were rehydrated in 50:50 PBSTx:MeOH for 10 min at RT, followed by a further 10 min incubation in PBSTx. Worms were then incubated in 1 ml bleaching solution (1.77 ml H2O, 100 µl formamide, 50 µl 20X SSC, 80 µl 30% hydrogen peroxide; all Sigma Aldrich) for 1 h under bright light [100]. The remainder of the protocol was carried out as described by King & Newmark [98] and is outlined below. A full protocol is available at https://www.protocols.io/view/fluorescent-in-situ-hybridisation-for-juvenile-fas-3byl4q7xjvo5/v1. Following the removal of bleaching solution, worms were rinsed in 1x SSC (20x SSC diluted to 1x in DEPC treated H2O) for 10 min before being rinsed in PBSTx for 10 min on rotation. To permeabilise the worms, they were subjected to a 15 min incubation in 1 ml of proteinase K solution (50 µl 10% SDS and 2.75 µl of 20 mg/ml proteinase K [Sigma Aldrich] in 4.95 ml PBSTx) prior to being postfixed in 4% formaldehyde for 10 min on rotation. Worms were then washed for a further 10 min in PBSTx, on rotation, before being transferred to 1:1 PBSTx:PreHybridisation solution (Prehyb: 25 ml de-ionized formamide, 12.5 ml 20X SSC, 100 µl of 50 mg/ml yeast RNA, 5 ml 10% (v/v) Tween-20 [Sigma Aldrich], and 7.4 ml DEPC-treated H2O) in 35 µm incubation baskets (Intavis) within an 18 well plate and incubated on a shaker at RT for 10 min. Baskets containing worms were then transferred to wells containing PreHybridisation solution before being placed in a hybridisation oven where they were incubated at 52°C for 2 h. Following this 2 h incubation, worms were hybridised by transferring baskets to wells containing Riboprobe mix (20 µl of 50 ng/µl target riboprobe was heated to 80°C for 5 min and placed on ice prior to the addition of 980 µl of hybridisation solution) and incubating overnight at 52°C in a Boekel Scientific Shake’N Bake™ hybridisation oven.

The following morning, 500 µl of Riboprobe mix was removed from wells and replaced with 500 µl of preheated 2X SSCx (20X SSC stock diluted to 2X in ddH2O + 0.1% Triton X-100) in which worms were incubated for 20 min. Baskets were then transferred to new wells containing 2X SSCx and incubated for 20 min. This was repeated an additional 2x. Worms then underwent 4x 20 min washes in 0.2X SSCx (20X SSC stock diluted to 0.2X in ddH2O + 0.1% Triton X-100). All SSC washes were carried out at 52°C. Following the final 0.2X SSCx wash, baskets were transferred to wells containing TNTx (1 L ddH2O, 12.11 g Tris Base, 8.77 g NaCl, 3 ml Triton X-100, pH 7.5) and incubated for 10 min x2 on a shaker at RT before being transferred to wells containing blocking solution (500 µl horse serum and 500 µl Roche Western Blocking Reagent [both Sigma Aldrich] diluted in 9 ml TNTx), in which they were incubated at RT for 1 h. Baskets were then transferred to wells containing antibody solution (anti-DIG-POD [Sigma Aldrich] diluted in blocking solution, 1:2000), and incubated overnight at 4°C.

Worms were washed in TNTx for 5 min, 10 min and 6x for 20 min at RT via the transfer of baskets to new wells prior to a 10 min incubation in freshly made tyramide solution (1 ml tyramide signal amplification (TSA) buffer [2 M NaCl; 0.1 M Boric acid, pH 8.5], 6 µl 5% H2O2, 1 µl 4 IPBA [20 mg 4-iodophenylboronic acid in 1 ml dimethylformamide (DMF)], 2 µl TAMRA). The fluor-conjugated NHS ester employed in the preparation of tyramide conjugates was TAMRA (5-(and-6)-carboxytetramethylrhodamine, Sigma Aldrich).

During tyramide incubation and for the remainder of the protocol, plates were protected from light. Post tyramide incubation, EdU detection and DAPI staining were performed as detailed below. During all wash/incubation steps post SSCx washes, plates containing baskets of worms were placed on a shaker at 100 rpm. Worms were mounted in 10 µl Vectashield (Vector Laboratories) on standard microscope slides and coverslips were sealed with clear nail polish. Only those RNAi targets yielding phenotypes were subjected to FISH.

### RNA interference (RNAi)

Double-stranded (ds)RNA templates specific for each target gene were amplified from juvenile *F. hepatica* cDNA using primers labelled with the T7 promoter sequence; 5’-TAATACGACTCACTATAGGGT-3’. Primers were designed using Primer3Plus and amplicons were later sequence confirmed by Eurofins Genomics (https://eurofinsgenomics.eu/en/). Primer sequences used to generate dsRNA templates can be found in S1 Table. Amplicons were generated using end-point PCRs (as above) and sizes confirmed on a 2 % agarose gel. dsRNA templates were purified using the ChargeSwitch™ PCR Clean-Up Kit (#CS12000, ThermoFisher Scientific) and dsRNA synthesis performed using the T7 RiboMAX™ Express RNA System (#P1700, Promega) according to manufacturer’s instructions. The resulting dsRNA constructs were re-suspended in nuclease free H2O and their RNA concentrations and purities measured using a DeNovix DS-11 FX spectrophotometer/fluorometer. All dsRNAs were stored at −20 °C as single use aliquots.

All RNAi experiments were performed in triplicate. Worms were soaked in 100 ng/µl of target (or control) dsRNA in 50 µl RPMI reactions for 24 h under standard culture conditions. dsRNA exposures were repeated biweekly for four weeks from the date of excystment. Between exposures, worms were returned to CS50 and maintained as standard. All RNAi experimental setups included an untreated control group (no dsRNA), while an exogenously derived dsRNA template (bacterial neomycin phosphotransferase [U55762]) served as a negative control. Upon completion of trials, juveniles were transferred to 2 ml round-bottomed Eppendorf tubes and snap frozen in liquid nitrogen prior to messenger (m)RNA extraction and reverse transcription.

### Quantification of RNAi target transcript knockdown

Frozen samples from each treatment group were lysed using a Qiagen TissueLyser LT at 50 oscillations/minute for one minute. Poly-adenylated mRNA was then extracted from each sample using the Dynabeads™ mRNA Direct™ Kit (#61012, ThermoFisher Scientific) before being treated with DNase (Turbo DNA-*free*™ Kit, #AM1907, ThermoFisher Scientific) and reverse transcribed to cDNA (High-Capacity RNA-to-cDNA™ Kit, #4387406, ThermoFisher Scientific). All cDNAs were diluted 1:1 in nuclease-free H2O prior to use. Target transcripts were amplified via qPCR on a Qiagen RotorGene Q 5-plex HRM instrument using the following cycling parameters: 95°C for 10 min, followed by 40 cycles of 95°C 10 s, 60°C 15 s and 72°C for 30 s. qPCRs were performed using 10 µl reactions consisting of 5 μl SensiFAST SBYR No-ROX Kit (#BIO-98005, Bioline), 0.2 μM of each primer and 2 μl of relevant cDNA (or H2O in the case of no template controls). All reactions were performed in triplicate, including no-template controls. *F. hepatica* glyceraldehyde 3-phosphate dehydrogenase (*Fh*GAPDH, [AY005475]) served as a reference gene. Melt-curve analyses were employed as standard.

Pfaffl’s augmented ΔΔCt method [101] was employed to calculate relative gene expression. Ratios of target transcript abundance relative to the untreated control were then converted to percentage transcript expression where 100% represents no change. Transcript expression was plotted for both target and control-dsRNA-treated groups relative to no dsRNA control.

### Compound exposures

Pyrvinium pamoate (#HY-A0293, MedChem Express), FZM1 (1-(3-Hydroxy-5-(thiophen-2-yl)phenyl)-3-(naphthalen-2-yl)urea, #5343580001, Sigma Aldrich) and triclabendazole (TCBZ; #1681611, Sigma Aldrich) were dissolved to the desired concentrations using dimethyl sulphoxide (DMSO, #D2650, Sigma Aldrich) within NoStick® hydrophobic microtubes. All compound exposures were carried out in 3 ml reactions within 35×10 mm Petri dishes (#82.1135.500, Sarstedt), with compounds being added at 1:1000, such that the final concentration of DMSO was 0.1% (v/v). All assays were performed alongside TCBZ-treated and vehicle (DMSO)-treated positive and negative control groups, respectively. Parasites were exposed to compounds for 18-hours biweekly from the date of excystment for three weeks under standard culture conditions. Prior to exposures, worms were washed in RPMI to remove excess chicken serum. Phenotypic observations were recorded immediately after the second exposure each week. Post compound exposures, worms were washed in RPMI to remove any residual drug before being returned to CS50 and cultured as standard. Following the final exposure, worms from treatment groups exhibiting growth phenotypes were incubated in EdU for 24 h to allow for the visualisation of any effects on neoblast-like cell proliferation.

### Labelling neoblast-like cells using 5-ethynyl-2′-deoxyuridine (EdU)

Following the final dsRNA exposure on day 28 of RNAi experiments, or after final compound exposures, a cohort of worms from control groups and each treatment group producing a growth phenotype were incubated in 500 µM EdU (dissolved in 1x PBS) in CS50 under standard culture conditions for 24 h. Worms were then washed in RPMI and transferred to a 20 µl droplet of 4% paraformaldehyde (PFA) in PBS within a Petri dish and flat fixed under a weighted coverslip for 10 min. Following removal of the coverslip, worms were collected into NoStick® hydrophobic microtubes containing 1 ml 4% PFA, and free fixed O/N at 4°C, while under constant rotation.

EdU detection was performed using a modified version of an *S. mansoni* EdU staining protocol [102]. Fixed, EdU-labelled worms were permeabilised for 30 min in PBSTx (PBS containing 0.5% Triton-X-100). EdU detection was performed via a 30-min incubation in a 6-carboxyfluorescein (6-FAM) azide conjugate (Metabion) solution while protected from light. Worms were then washed 2x in PBSTx and counterstained via a 20-min incubation in 1:1000 4′,6-diamidino-2-phenylindole (DAPI, 1 mg/ml, Sigma Aldrich) in PBS, again, while protected from light. Two final 5-min washes were performed in PBSTx prior to mounting (as previously described). All steps were performed under constant rotation at RT. A full version of the protocol is available at https://www.protocols.io/private/41904A32719811EE95C60A58A9FEAC02.

### Immunocytochemistry (ICC)

Following the final dsRNA exposure on day 28 of *Fh*CTNNB1 RNAi experiments, a cohort of worms from each treatment group were processed for fixation (as above). ICC was performed using neuropeptide F (NPF) antiserum (raised against the C-terminal decapeptide-amide of NPF from *Moniezia expansa* [103]) and tetramethylrhodamine-isothiocyanate (TRITC)-conjugated phalloidin (#P1951, Sigma Aldrich) to enable the visualisation of a significant portion of the peptidergic nervous system and musculature, respectively. Worms were incubated in rabbit anti-NPF primary antiserum (1:1000) for 72 h, washed in antibody diluent (AbD; 0.1 M PBS, 0.1% (v/v) Triton X-100, 0.1% (w/v) bovine serum albumin) and incubated for a further 48 h in 1:100 fluorescein isothiocyanate (FITC)-labelled anti-rabbit secondary antiserum (Sigma Aldrich). A final overnight incubation in 1:100 TRITC-labelled phalloidin (200 ng/ml, Sigma Aldrich) was performed before three 15-min washes in AbD prior to mounting (as above). The dilution of antisera and phalloidin was achieved using AbD and all incubation periods were carried out under constant rotation at 4°C.

### Phenotypic analyses

Parasite motility and size were recorded following the second dsRNA/compound exposure each week for the duration of RNAi and compound assays, respectively. Darkfield videos, each one minute in length, were captured using an Olympus SZX10 microscope with attached Olympus SC50 camera. Videos were analysed using the wrMTrck plugin for ImageJ (https://www.phage.dk/plugins/wrmtrck.html). Changes in worm length between frames provided values for individual length changes (µm/minute) and served as a measure of motility. The motility of individual worms was then normalised to the average movement of untreated (no-dsRNA) and vehicle (DMSO) controls for RNAi and compound assays, respectively. Values were presented as percentage motility relative to the relevant control. Area values (µm^2^) for individual worms were also obtained using wrMTrck.

At the conclusion of experiments, brightfield images of each treatment group were captured using a Leica M205 C stereomicroscope with attached Leica MC190 HD camera to document the presence of any aberrant, morphological changes following RNAi and compound exposures. Neoblast-like cell activity was quantified via the enumeration of EdU^+^ nuclei using the cell counter plugin for ImageJ (https://imagej.nih.gov/ij/plugins/cell-counter.html) which were then normalised to worm area, giving values per 1000 µm^2^.

### Confocal microscopy

All FISH, EdU-labelled and ICC specimens were imaged using a Leica TCS SP8 inverted confocal scanning laser microscope. Images were captured as maximally projected z-stacks, each generated from 30-40 optical sections between the dorsal and ventral surfaces of whole worms.

### Statistical Analyses

All statistical analyses were performed and graphs produced using GraphPad Prism 9 (La Jolla California USA, www.graphpad.com). Each dataset was first tested for normality. Where data were normally distributed, parametric tests including ANOVAs and t-tests were performed, while the nonparametric Kruskal-Wallis and Mann-Whitney tests were employed where data were not normally distributed. Post-hoc analyses were carried out using Dunn’s, Dunnett’s and Tukey’s multiple comparisons tests to identify differences between several groups.

## ACKNOWLEDGEMENTS

Funding for this study was provided by the Biotechnology and Biological Sciences Research Council and Boehringer Ingelheim Ltd. (BB/T002727/1) and the Department of Agriculture, Environment and Rural Affairs.

## Supplementary

**S1 Figure.**
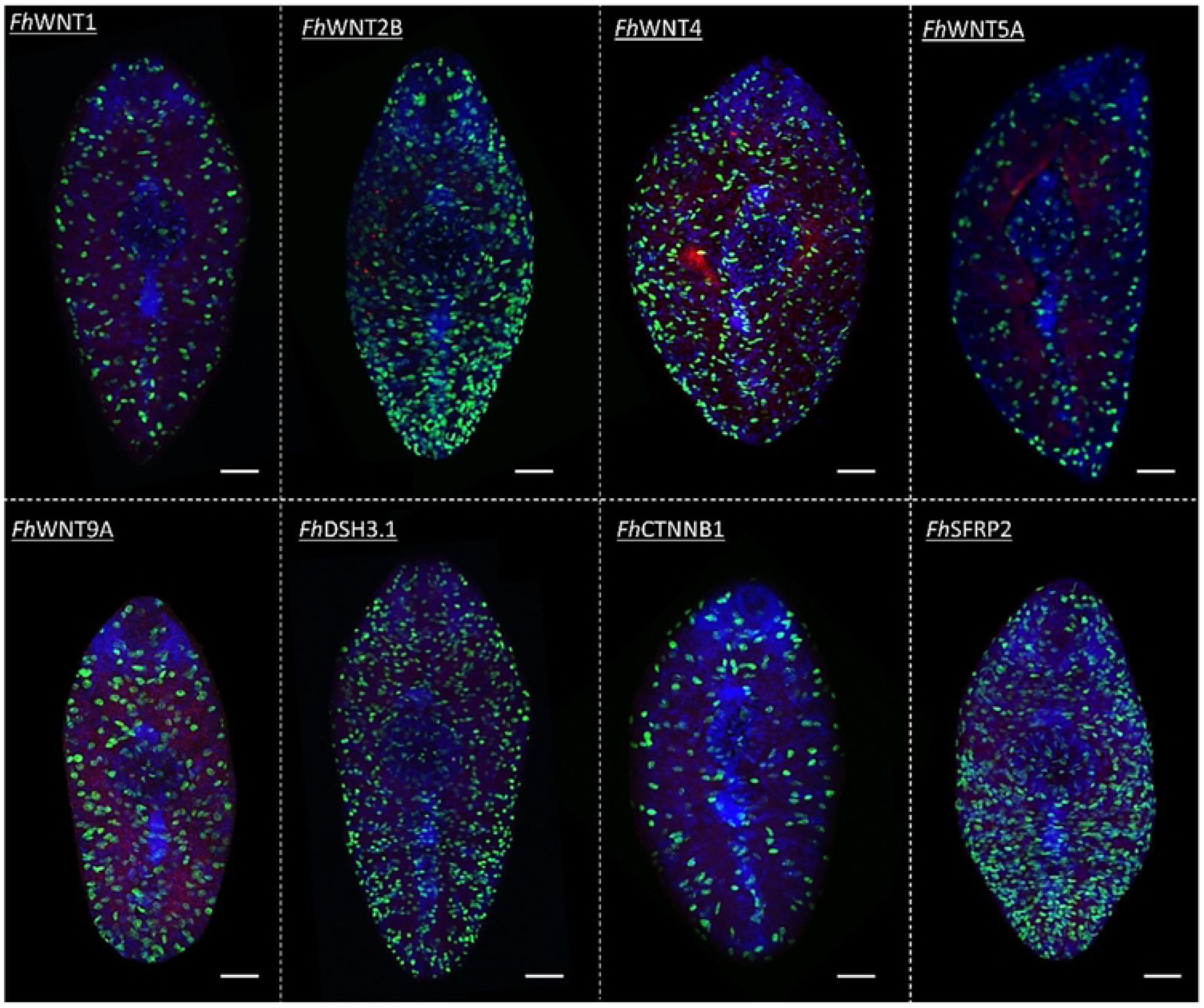
Negative controls demonstrate the specificity of Fh\Nnt pathway component transcript localisation. Fluorescence in situ hybridisation (FISH) negative control juvenile Fasciola hepatica exposed to sense (forward) strand RNA probes. Red (TAMRA) indicates non-specific binding, green fluorescence denotes EdU+ (neoblast-like) cells. DARI (blue) served as a counterstain. Scale = 50 pm.

**S2 Figure.**
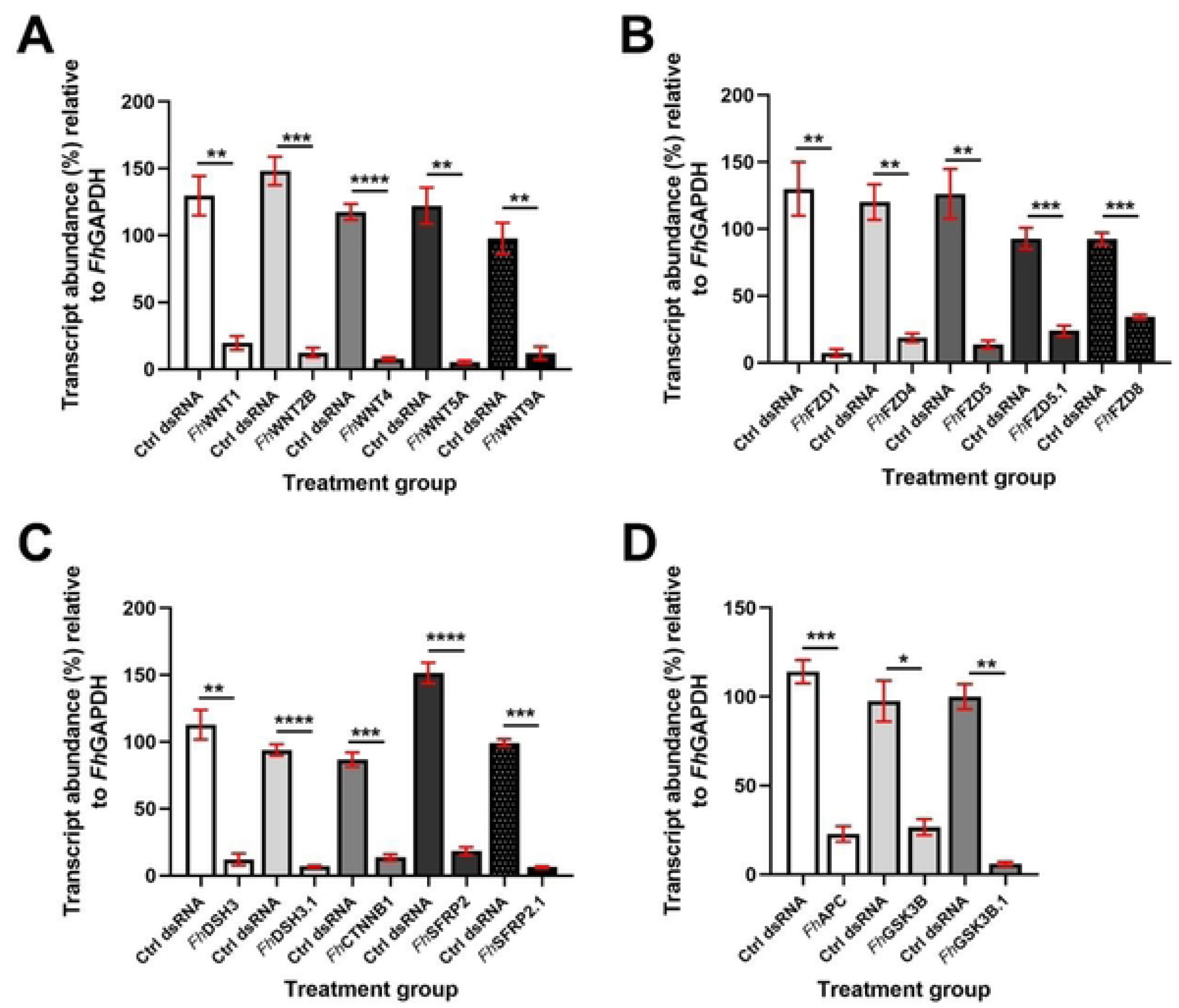
Juvenile liver fluke Wnt/p-catenin pathway component gene transcripts can be silenced using RNA interference. Transcript knockdown following RNA interference of Wnt/p-catenin pathway components in 4-week-old juvenile Fasciola hepatica (A) *Fh*Wnts (B) *Fh*FZDs (C) Other active Wnt signalling targets; *Fh*DSH, *Fh*CTNNB1 and *Fh*SFRP (D) Destruction complex targets (n=3). Data show mean expression (±SEM) of target transcript in control and target dsRNA-treated juveniles relative to untreated controls, using *Fh*GAPDH as a housekeeping gene. Transcript knockdown was measured following nine 24-hour dsRNA exposures over a period of four weeks. Statistical analyses were performed using unpaired t-tests. *, p<0.05; **, p<0.01; ***, p<0.001; ****, p<0.0001.

**S3 Figure.**
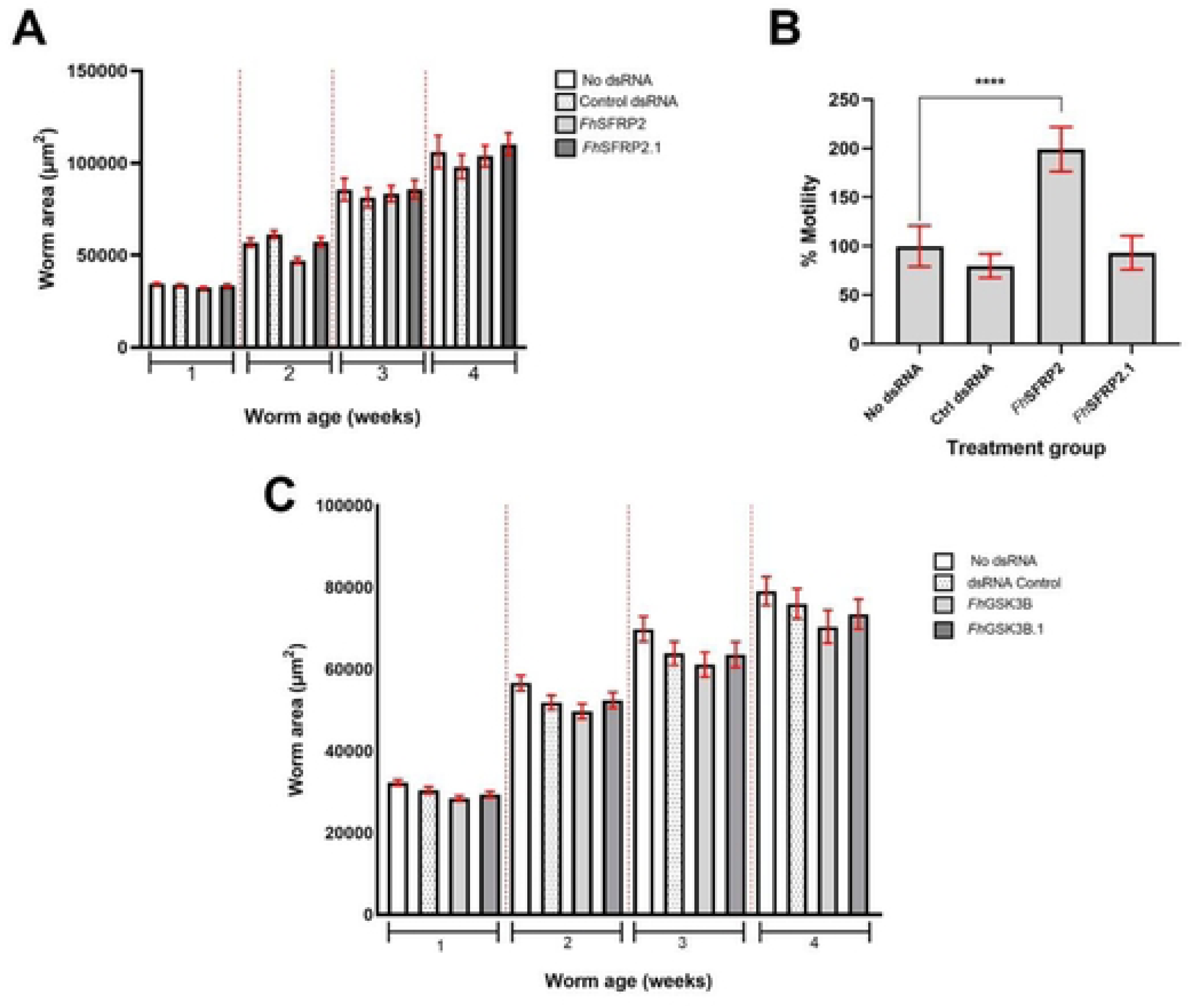
The silencing of selected Wnt/p-catenin pathway components known to inhibit Wnt signalling does not impact juvenile liver fluke growth. A) Growth of juvenile Fasciola hepatica following RNAi-mediated silencing of FfrSFRPs. B) Motility analysis of four-week old *Fh*SFRP-silenced juveniles. C) Growth of juvenile F hepatica following RNAi-mediated silencing of F/X3SK3B. Worm area measured in pm^2^, data presented as pm^2^±SEM. Statistical analyses were performed using Kruskal Wallis with Dunn’s post hoc tests. ****, p<0.0001.

---

**S1 Table.**
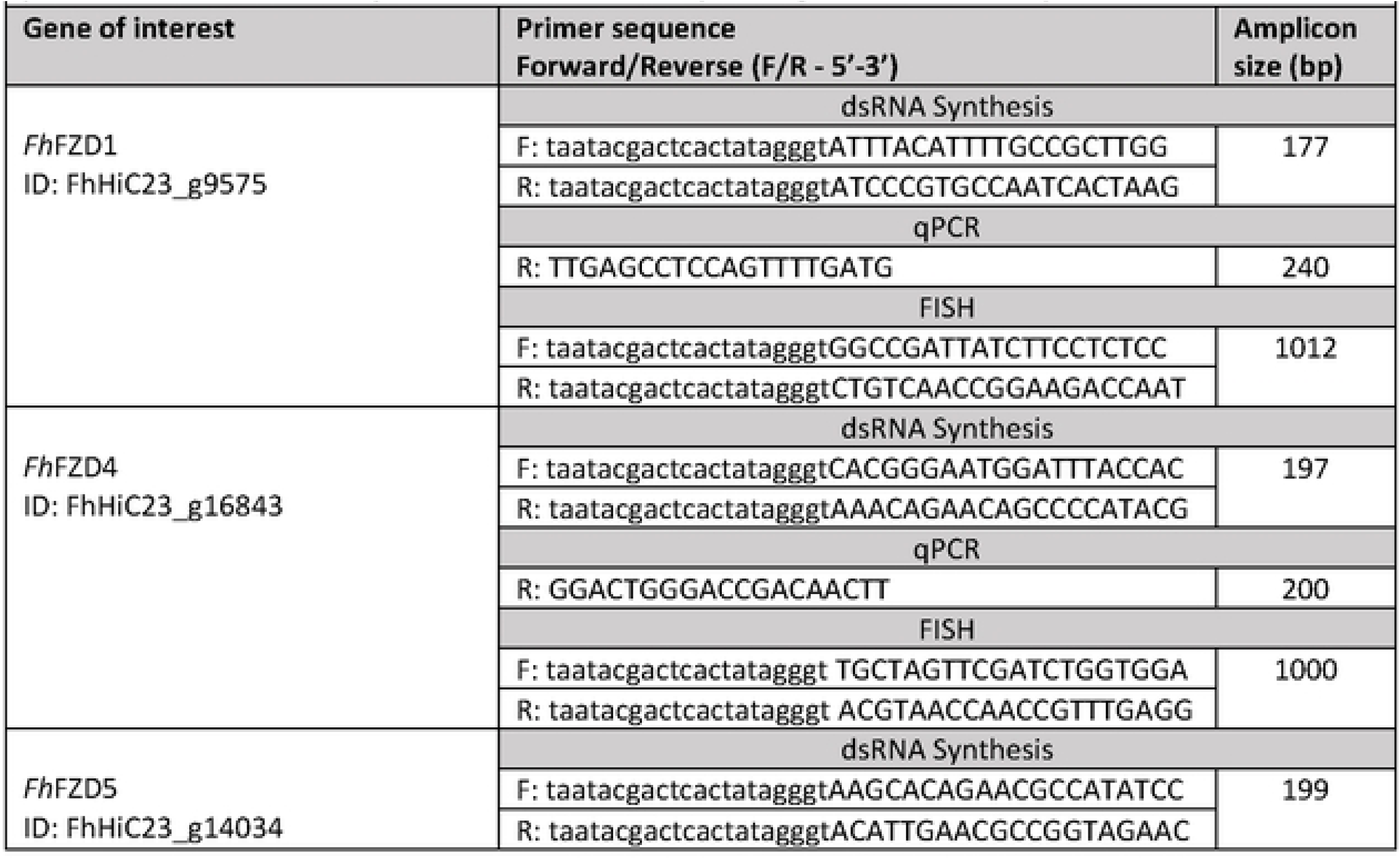
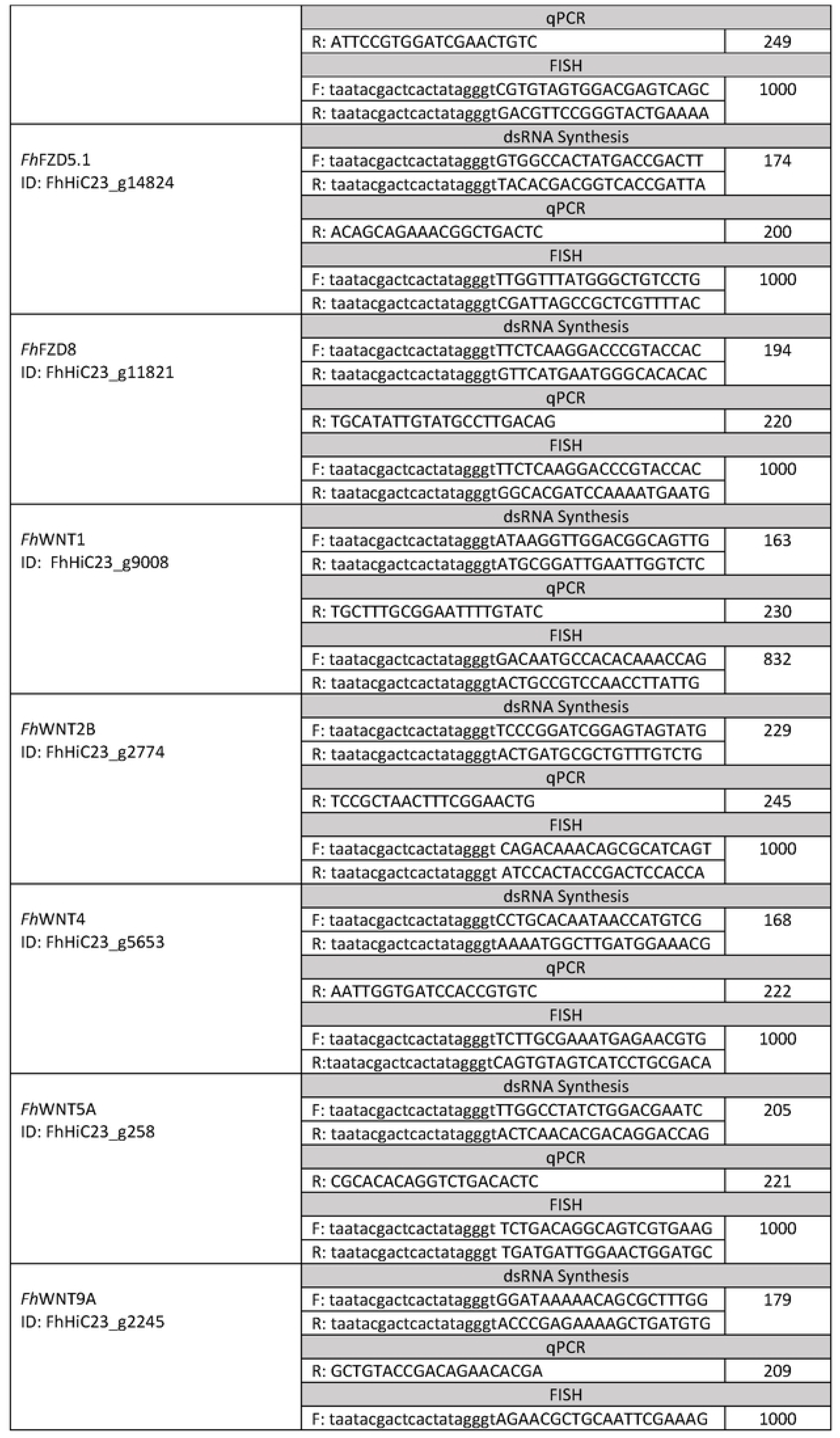
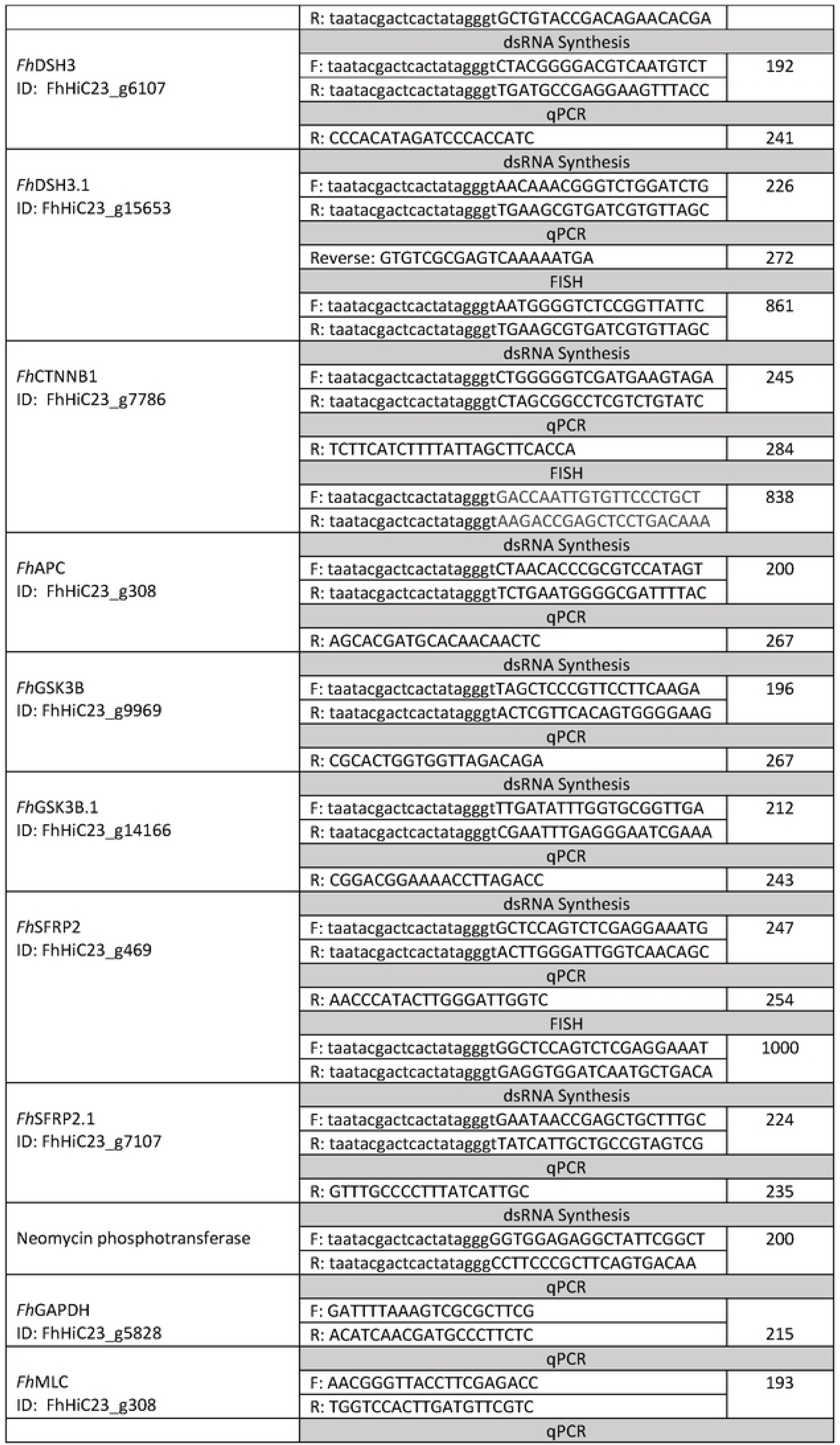
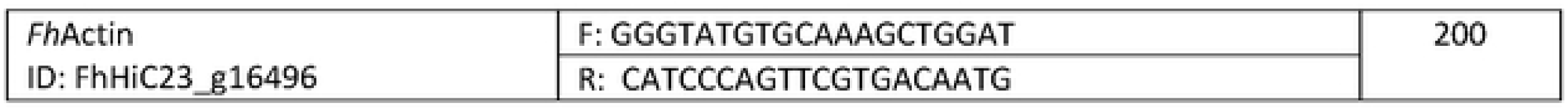
Oligonucleotide primers used in double stranded (ds)RNA synthesis, qPCR analyses and fluorescence in situ hybridisation (FISH) of FhWnt pathway targets. N.B. qPCR reverse primers were used in conjunction with the corresponding dsRNA forward primer.

**S2 Table.**
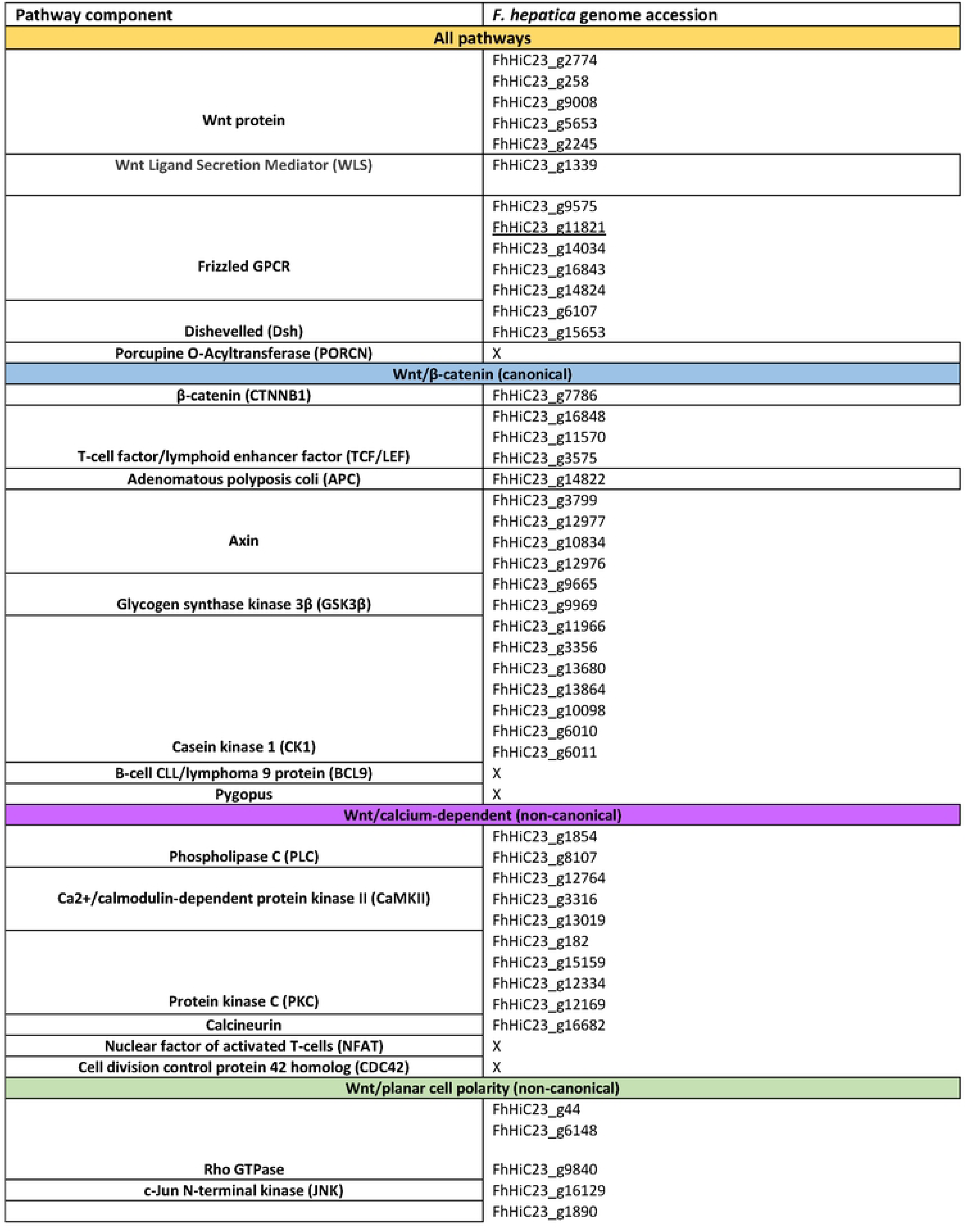
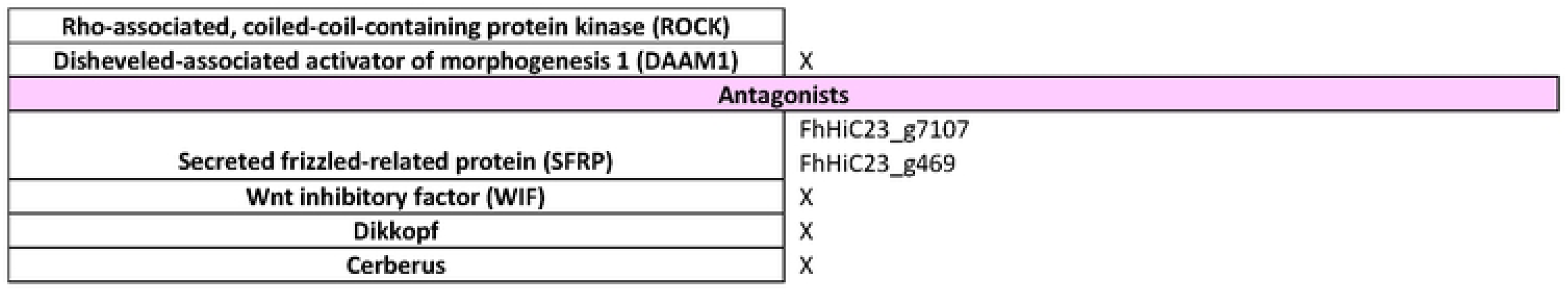
Fasciola hepatica \Nr\t signalling pathway component gene IDs. Those pathway components lacking homologues in F hepatica are denoted by an ‘X’.

**S3 Table.** Fasciola hepatica Wnt pathway component RNAi targets and their orthologues (top BLAST hits) in higher and related species.

| *F. hepatica* genome accession | Top hit: *H. sapiens* (taxid: 9606) | Top hit: *M. musculus* (taxid: 10090) | Top hit: *S. mediterranea* (taxid: 79327) | Top hit: *D. melanogaster* (taxid: 7227) | Top hit: *S. mansoni* (taxid: 6183) | Top hit: *E. multilocularis* (taxid: 6211) |
| --- | --- | --- | --- | --- | --- | --- |
| FhHiC23_g9008 | Proto-oncogene Wnt-1 | Protein Wnt-10A | Protein Wnt-1 | Wingless | Smp_248080 | EmuJ_000349900 |
| FhHiC23_g2774 | Protein Wnt-2b | Protein Wnt-10B | Protein Wnt-2-1 | Wingless | Smp_167140 | EmuJ_000748600 |
| FhHiC23_g5653 | Protein Wnt-4 | Protein Wnt-4 | Protein Wnt-a | Wnt-2 protein | Smp_332550 | EmuJ_000211300 |
| FhHiC23_g258 | Protein Wnt-5a | Protein Wnt-5B | Protein Wnt-5 | Wnt-2 protein | Smp_145140 | EmuJ_000804000 |
| FhHiC23_g2245 | Protein Wnt-9a | Protein Wnt-9a | Protein Wnt-11-1 | Protein Wnt-5 | Smp_156540 | EmuJ_000907500 |
| FhHiC23_g9575 | Frizzled-1 | Frizzled-1 | Frizzled-1 | Frizzled class receptor | frizzled class receptor 1 | EmuJ_000682100 |
| FhHiC23_g16843 | Frizzled-4 | Frizzled-4 | Frizzled-4 | Frizzled-4 | Smp_247930 | EmuJ_000085700 |
| FhHiC23_g14034 | Frizzled-5 | Frizzled-8 | Frizzled 5/8-3 | FBgn0001085 | Smp_155340 | EmuJ_000996400 |
| FhHiC23_g14824 | Frizzled-5 | Frizzled-8 | Frizzled 5/8-4 | Frizzled-2 | Smp_139180 | EmuJ_000996400 |
| FhHiC23_g11821 | Frizzled-8 | Frizzled-8 | Frizzled-4 | Frizzled-2 | Smp_174350 | EmuJ_000438200 |
| FhHiC23_g6107 | Dishevelled segment polarity protein 3 | Dishevelled segment polarity protein 3 | Dishevelled like protein, DVL-b | Dishevelled segment polarity protein | Smp_020300 | EmuJ_000118000 |
| FhHiC23_g15653 | Dishevelled segment polarity protein 3 | Dishevelled segment polarity protein 3 | Dishevelled like protein, DVL-b | Dishevelled segment polarity protein | Smp_347960 | EmuJ_000423700 |
| FhHiC23_g7786 | Catenin Beta 1 | Catenin Beta 1 | Catenin Beta 1 | Armadillo | Smp_023550 | EmuJ_001007700 |
| FhHiC23_g9969 | Glycogen synthase kinase 3-beta | Glycogen synthase kinase 3-beta | Glycogen synthase kinase 3.1 | Gasket | Smp_008260 | EmuJ_000352700 |
| FhHiC23_g14166 | Glycogen synthase kinase 3-beta | Glycogen synthase kinase 3-beta | Glycogen synthase kinase 3.1 | Shaggy | Smp_008260 | EmuJ_000980300 |
| FhHiC23_g7107 | Secreted Frizzled-related protein 2 | Secreted Frizzled-related protein 2 | Secreted Frizzled-related protein 1 | Frizzled gene exon 1 | Smp_068680 | EmuJ_000838700 |
| FhHiC23_g469 | Secreted Frizzled-related protein 2 | Secreted Frizzled-related protein 2 | Secreted Frizzled-related protein 1 | Frizzled gene exon 1 | Smp_062560 | EmuJ_000838700 |
| FhHiC23_g14822 | Adenomatous polyposis coli | Chain A, Adenomatous polyposis coli protein | Adenomatous polyposis coli-like protein | Adenomatous polyposis coli | Smp_343270 | EmuJ_001070700 |

